# Driving proteomic imbalance in malignancy provokes proteomic catastrophe and confers tumor suppression

**DOI:** 10.1101/2024.05.24.595838

**Authors:** Babul Moni Ram, Omprakash Shriwas, Meng Xu, Kun-Han Chuang, Chengkai Dai

**Author notes:** School of Basic Medical Sciences, Fujian Medical University, Fuzhou, Fujian 350108, China (Kun-Han Chuang).

## Abstract

Unlike genomic instability, the role of proteomic instability in cancer remains poorly defined. Heat shock factor 1 (HSF1), a master regulator of the proteotoxic stress response, preserves proteome integrity under stress and is increasingly recognized as an oncogenic enabler. In Neurofibromatosis type I (*NF1*)-deficient malignant peripheral nerve sheath tumor (MPNST) cells, *HSF1* loss induces widespread protein polyubiquitination, aggregation, and tumor-suppressive amyloidogenesis, yet is dispensable in non-transformed Schwann cells. Mechanistically, HSF1 protects the mitochondrial chaperone HSP60 from toxic soluble amyloid oligomers. To adapt to compromised protein quality, *HSF1*-deficient MPNST cells activate JNK to repress mTORC1-dependent translation, reducing protein load. Stimulating mTORC1 in these *HSF1*-deficient cells drives catastrophic proteomic imbalance, triggering pronounced cell death, in part through unchecked amyloidogenesis, and suppressing tumor growth *in vivo*. Thus, HSF1 safeguards the cancer proteome, enabling the oncogenic capacity of mTORC1. This proof-of-principle study establishes the induction of proteomic catastrophe as a new paradigm for combating malignancy.

## INTRODUCTION

Compared to genomic instability, proteomic instability in cancer is underappreciated. Malignant transformation triggers proteotoxic stress and destabilizes the proteome.^1, 2^ Multiple mechanisms, including reactive oxygen species, genetic mutations, and dysregulated protein translation, contribute to increased protein damage, misfolding, and aggregation.^1–3^ Notably, amyloidogenesis, a hallmark of neurodegenerative disorders,^4^ also occurs in cancer.^5^ Unlike neurons, however, cancer cells can contain amyloid formation and mitigate its cytotoxic effects. A key component of this defense is heat shock factor 1 (HSF1), which drives the heat shock response/proteotoxic stress response (HSR/PSR), a potent cytoprotective transcriptional program.^6–8^ Consistent with its role in repressing proteomic instability, HSF1 has emerged as a powerful oncogenic enabler independent of driver mutations.^9–15^

In contrast to its dispensability in normal and non-transformed cells, HSF1 is essential in many cancer cell lines,^9,13,16–18^ due to elevated intrinsic proteotoxic stress. This dependency, termed “non-oncogene addiction” to HSF1,^2,19^ highlights its potential as a therapeutic target.

Neurofibromatosis type 1 (NF1) is a cancer predisposition syndrome affecting approximately 1 in 3000 individuals worldwide.^20^ About 30-50% of NF1 patients develop benign plexiform neurofibromas, which can progress into highly aggressive malignant peripheral nerve sheath tumors (MPNSTs) with poor prognosis.^21^ The *NF1* gene encodes neurofibromin, a 250-280 kDa RAS GTPase activating protein (GAP) that negatively regulates RAS signaling.^22^ Loss of *NF1* in MPNST cells leads to hyperactive RAS/MAPK signaling, which not only promotes cellular growth but also drives constitutive HSF1 activation.^13^ Accordingly, HSF1 overexpression and activation are observed in both experimental MPNST models and primary NF1 tumors.^13^ Significantly, *Hsf1* deficiency impaired NF1-associated tumorigenesis in mice, partly through attenuation of oncogenic RAS signaling.^13^

Utilizing the NF1 tumor model system, we herein identify HSF1 as a central regulator that counteracts proteomic instability in cancer. We demonstrate that amyloidogenesis occurs in *NF1*-deficient MPNST cells, driven in part by hyperactive mTORC1 signaling and increased protein translation. Inhibition of HSF1 destabilizes the cancer proteome, leading to widespread protein polyubiquitination, aggregation, and amyloid formation. This proteomic instability, ultimately, causes cytotoxicity. To survive HSF1 inhibition, MPNST cells activate JNK to suppress mTORC1 and reduce protein synthesis. Disruption of this adaptative response via JNK inhibition exacerbates amyloidogenesis and cell death. Similarly, mTORC1 stimulation—via genetic *TSC2* depletion or supplementation with leucine or its analog—synergizes with HSF1 inhibition to induce profound proteomic chaos and widespread non-apoptotic cell death. Notably, blocking amyloidogenesis significantly rescues cell viability. Together, these findings reveal a critical role for HSF1 in antagonizing proteomic instability in cancer cells, representing a general pro-oncogenic mechanism. This work provides proof of principle that inducing proteomic catastrophe may serve as a next-generation anti-cancer therapeutic strategy.

## RESULTS

### HSF1 counters proteomic instability in *NF1*-deficient MPNST cells

We previously demonstrated that *NF1* loss leads to constitutive HSF1 activation and that *Hsf1* deficiency impairs tumorigenesis in *Nf1*^+/-^; *p53*^+/-^ (NPcis) mice.^13^ Consistent with these findings, analysis of human cancers revealed that *NF1* deletion is associated with elevated *HSF1* gene expression (Figure 1A), and *NF1* and *HSF1* expression levels are inversely correlated (Figure S1A). Importantly, among patients with *NF1*-deleted tumors, higher *HSF1* expression correlates with reduced survival (Figure 1B). Together, these data support a pro-oncogenic role for HSF1 in *NF1*-deficient malignancies.

**Figure 1:**
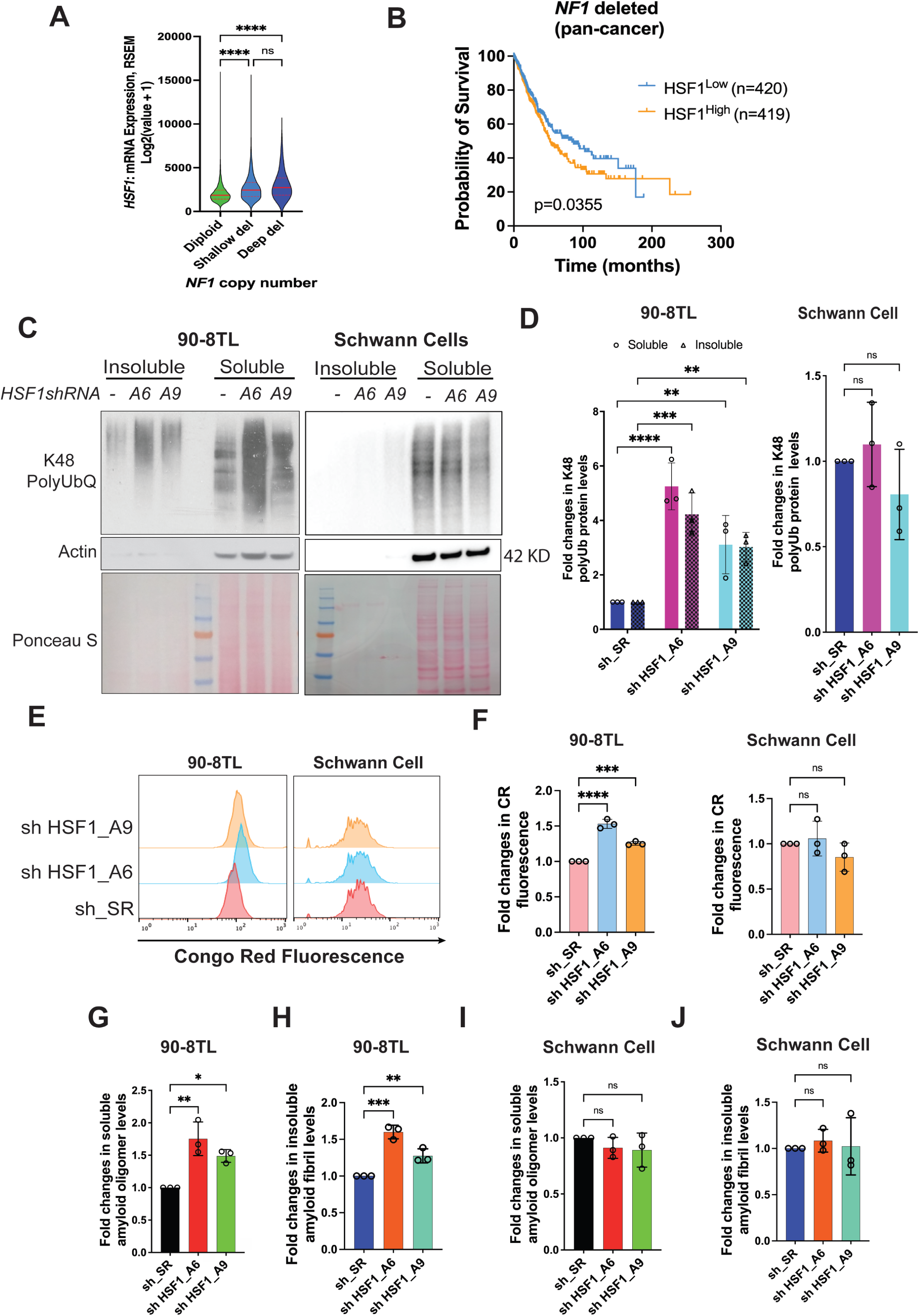
HSF1 is required to suppress proteomic instability and amyloidogenesis in MPNST cells. (A) Violin plots of *HSF1* expression in human cancer tissues showing *NF1* deletion (median and IQR, n=6424, 1598, or 88, Kruskal-Wallis test). Data are generated by the TGCA Research Network (https://www.cancer.gov/tcga). (B) Kaplan-Meier survival curves of patients whose tumors harbor *NF1* deletion (n=420 or 419, Log-rank test). Data are generated by the TGCA Research Network (https://www.cancer.gov/tcga). (C) and (D) The impacts of *HSF1* depletion on protein polyubiquitination in both 90-8TL cells and primary human Schwann cells. Cells were transduced with lentiviral shRNAs for 4 days and global protein Lys48-specific ubiquitination was detected by immunoblotting in both detergent-soluble and -insoluble fractions of whole cell lysates (C). Protein ubiquitination was quantitated using Fiji imaging software (D) (mean ± SD, n=3 independent experiments, Two-way ANOVA). (E) and (F) Congo Red (CR) staining of 90-8TL cells and primary human Schwann cells with and without *HSF1* depletion, analyzed by flow cytometry (E). Quantitation of CR staining using the median fluorescence intensity (F) (mean ± SD, n=3 independent experiments, One-way ANOVA). (G)-(J) Quantitation of soluble amyloid oligomers and insoluble amyloid fibrils by ELISA using A11 and OC antibodies, respectively (mean ± SD, n=3 independent experiments, One-way ANOVA).

Given the critical role of HSF1 in protein homeostasis, we examined how its depletion impacts the proteome of *NF1*-deficient MPNST cells compared to *NF1*-proficient, non-transformed human Schwann cells (HSCs), both primary and immortalized. HSCs are the presumed cell of origin for MPNSTs.^20,22^ In three different MPNST cell lines (90-8TL, S462, and SNF96.2), transient *HSF1* depletion, using two previously validated lentiviral shRNAs,^9,13^ led to a marked increase in global protein Lys48-specific polyubiquitination, a molecular mark for proteasomal degradation,^23^ in both detergent-soluble and -insoluble fractions (Figures 1C, 1D, S1B, and S1C), indicative of widespread protein misfolding and aggregation. In contrast, no such increase was observed in HSCs (Figures 1C and 1D). These results indicate that HSF1 is specifically required to maintain proteomic stability in malignant Schwann cells.

While most cellular protein aggregates are amorphous, a subset adopts highly ordered β-sheet-rich structures known as amyloids, which are classically associated with neurodegenerative disorders, especially Alzheimer’s disease.^24^ To determine whether *HSF1* depletion induces amyloidogenesis, we stained cells with Congo red (CR), a widely applied amyloid-binding fluorescent dye.^25^ MPNST cells exhibited stronger CR staining than HSCs at baseline (Figure 1E), suggesting higher amyloid burden in malignant cells. Notably, *HSF1* depletion further increased CR staining in MPNST cells but not in HSCs (Figures 1E, 1F, S1D, and S1E). Similar results were obtained using Thioflavin T staining (Figure S1F), another popular amyloid dye.^25^ We further validated amyloid accumulation using ELISA with two conformation-specific antibodies (A11 and OC), which recognize soluble amyloid oligomers (AOs) and insoluble amyloid fibrils (AFs), respectively.^26,27^ *HSF1* depletion significantly increased both A11^+^ AOs and OC^+^ AFs across all three MPNST cell lines (Figures 1G, 1H, S1G-S1J), whereas no significant changes were detected in HSCs (Figures 1I and 1J). Collectively, these findings indicate that HSF1 is essential for maintaining proteome integrity in malignant Schwann cells, particularly by suppressing amyloidogenesis, while being largely dispensable in non-transformed Schwann cells.

### Unchecked proteomic instability is cytotoxic

It has become evident that HSF1 acts as a generic oncogenic enabler; nevertheless, a universal underlying mechanism has yet to be fully defined.

A healthy proteome enables the translation of genotypes into phenotypes. Thus, we reasoned that preservation of proteomic stability represents a fundamental pro-oncogenic mechanism of HSF1. To test this, we first determined whether proteomic instability—particularly amyloidogenesis—is causally related to the cytotoxicity induced by *HSF1* depletion in cancer cells. To inhibit amyloid formation, we employed CR, which exerts anti-amyloid effects through its amyloid-binding activity.^5,28,29^ Consistent with the concept of “non-oncogene addiction” in cancer, *HSF1* depletion markedly impaired the viability of MPNST cells. Significantly, CR treatment partially rescued this impaired viability (Figures 2A, S2A, and S2B), indicating that amyloidogenesis contributes to the cytotoxic effects of HSF1 inhibition. In contrast, neither *HSF1* depletion nor CR treatment affected the viability of HSCs (Figure 2B), congruent with their lack of proteomic instability and amyloidogenesis. Together, these findings demonstrate that unchecked proteomic instability—particularly amyloidogenesis—is cytotoxic. Accordingly, HSF1 enables oncogenesis, at least in part, by constraining proteomic instability.

**Figure 2:**
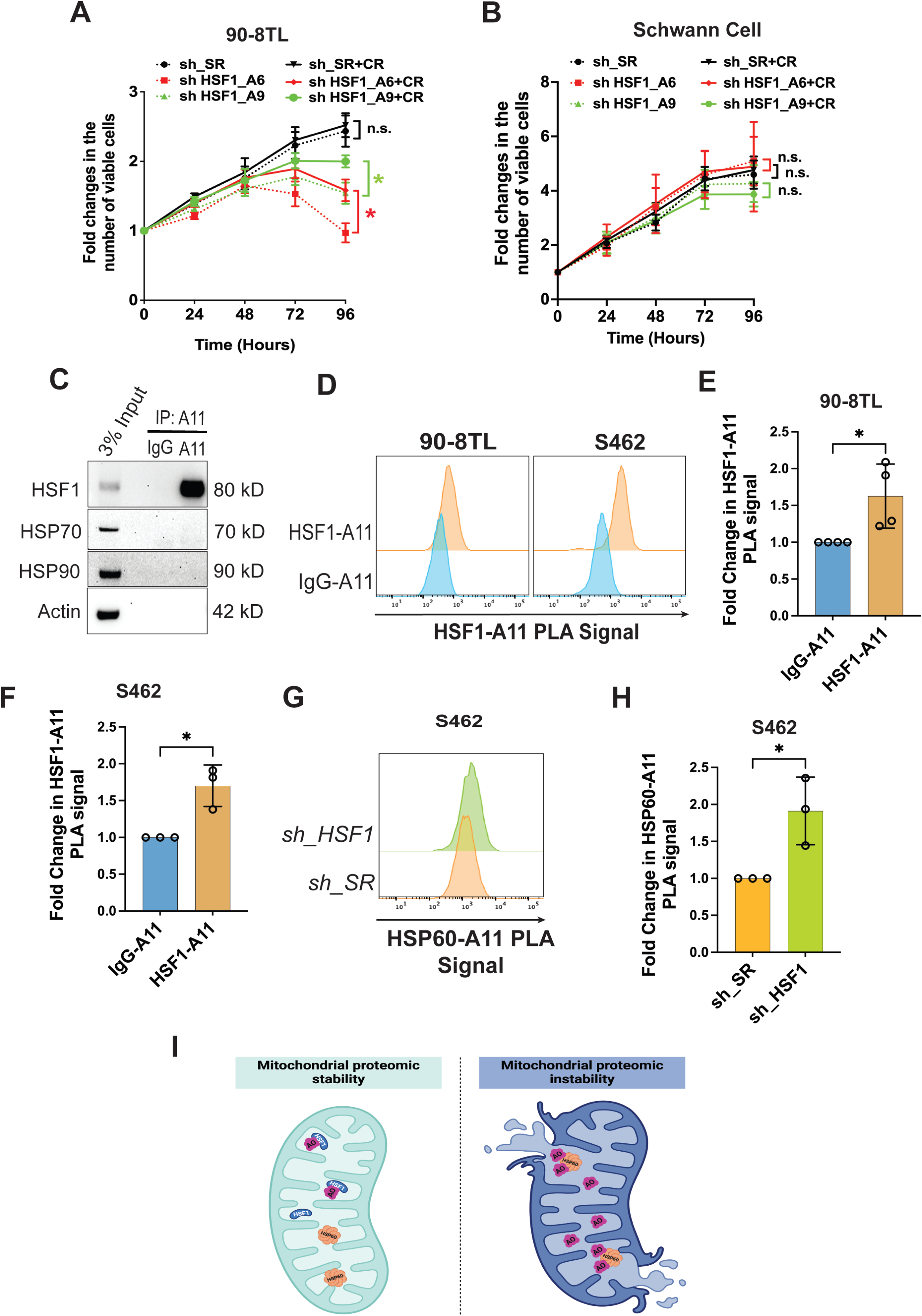
HSF1 counters the tumor-suppressive effects of amyloids in MPNST cells. (A) and (B) The growth curves of 90-8TL and primary human Schwann cells with and without *HSF1* depletion and 20 μM CR treatment (mean ± SD, n=3 independent experiments, Two-way ANOVA). (C) Co-IP of amyloid oligomers (AOs) with HSF1 in 90-8TL cells. (D)-(F) Detection of AO−HSF1 interactions by PLA flow cytometry in 90-8TL and S462 cells. Quantitation of the PLA signals using the median fluorescence intensity (mean ± SD, n=3 or 4 independent experiments, Student’s t test). (G) and (H) Detection of AO−HSP60 interactions by PLA flow cytometry in S462 cells following *HSF1* depletion (G). Quantitation of the PLA signals using the median fluorescence intensity (H) (mean ± SD, n=3 independent experiments, Student’s t test). (I) Schematic depiction of AO neuralization by HSF1 and HSP60 attack by AO following *HSF1* depletion in mitochondria.

### HSF1 neutralizes soluble amyloid oligomers to protect MPNST cells

How does HSF1 counter amyloid toxicity to enable the growth and survival of MPNST cells? In previous research using tissue overgrowth models, we discovered that HSF1, unexpectedly, physically neutralizes soluble AOs, thereby preventing their assault on HSP60.^30^ As an essential mitochondrial chaperone, HSP60 safeguards the mitochondrial proteome.^30–32^

We next asked whether this mechanism operates in cancer cells. Co-IP experiments indicated that A11^+^ AOs physically interacted with HSF1, but not major HSPs, in MPNST cells (Figure 2C), consistent with our previous findings.^30^ This AO–HSF1 interaction was further validated using an independent technique, *in situ* Proximity Ligation Assay (PLA) (Figures 2D-2F).

Significantly, *HSF1* depletion promoted the AO–HSP60 interaction (Figure 2G and 2H). Given that loss of *HSP60* destabilized the mitochondrial proteome,^30,32^ this AO–HSP60 interaction signifies the induction of mitochondrial damage and subsequent cytotoxicity. Collectively, our findings reveal that HSF1 protects MPNST cells, at least in part, by neutralizing soluble AOs, thereby preserving the mitochondrial chaperone HSP60 function (Figure 2I).

### MPNST cells react to HSF1 deficiency by activating JNK to attenuate translation

Given their dependency on HSF1, cancer cells are expected to engage adaptive responses to survive its inhibition. Previously, we uncovered that in *Hsf1*-deficient mice, the stress-responsive JNK kinase becomes constitutively activated, leading to disassembly of mTORC1 through phosphorylation of its core components, RAPTOR and mTOR.^33^ This inhibition of mTORC1 diminishes protein translation, leading to reduced cell size and whole-body lean mass in *Hsf1*-deficient mice.^33^

We therefore hypothesized that cancer cells might exploit this same molecular regulation for survival and adaptation. In MPNST cells, *HSF1* knockdown activated JNK, as indicated by increased phosphorylation at T183/Y185. This activation was accompanied by suppression of mTORC1 signaling, evidenced by diminished p70S6K T389 and 4EBP1 S65/T70 phosphorylation (Figures 3A, 3B, and S3A-S3D).^34,35^ In contrast, *HSF*1 knockdown had minimal effects on JNK and mTORC1 signaling in non-transformed HSCs (Figures 3A and 3B). Detected by PLA, in MPNST cells HSF1 interacted with JNK. Upon *HSF1* depletion, JNK–RAPTOR interactions increased, while mTOR–RAPTOR interactions decreased (Figures 3C-3E). Because the mTOR-RAPTOR interaction is essential for mTORC1 assembly and activity,^36^ these findings indicate that JNK senses HSF1 deficiency to repress mTORC1. Notably, this sensing mechanism appears more prominent in malignant cells than in non-transformed counterparts.

**Figure 3:**
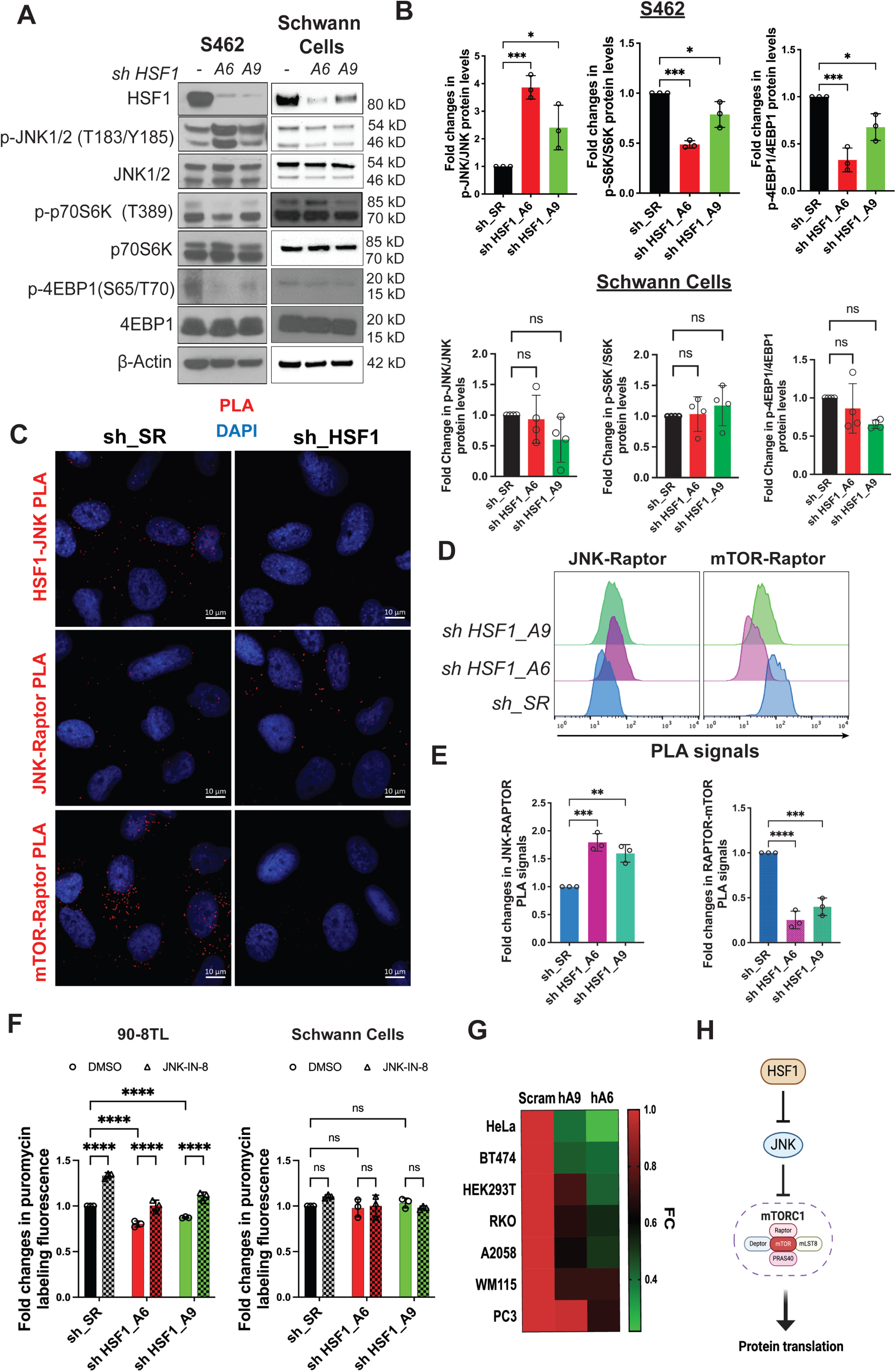
JNK senses *HSF1* deficiency to repress mTORC1 and protein translation in MPNST cells. (A) Detection of JNK activation and mTORC1 inhibition following *HSF1* depletion in S462 and primary human Schwann cells by immunoblotting. (B) Quantitation of (A) using Fiji imaging software (mean ± SD, n=3 independent experiments, One-way ANOVA). (C) Detection of HSF1-JNK, JNK-RAPTOR, and mTOR-RAPTOR interactions by PLA in S462 cells with and without *HSF1* depletion. (D) and (E) Detection and quantitation of JNK-RAPTOR and mTOR-RAPTOR interactions by PLA flow cytometry in 90-8TL cells (mean ± SD, n=3 independent experiments, One-way ANOVA). (F) Measurement of protein translation rates in 90-8TL and primary human Schwann cells with and without *HSF1* depletion by puromycin labeling, analyzed by flow cytometry. Cells were treated with 1 μM JNK-IN-8 for 24 hrs before labeling with 100 nM 6-FAM-dc-puromycin for 30 min (mean ± SD, n=3 independent experiments, Two-way ANOVA). (G) Heatmap of protein translation rates measured by puromycin labeling in diverse human cancer cell lines with and without *HSF1* depletion. (H) Schematic depiction of regulation of protein translation by HSF1.

Consistent with mTORC1 inhibition, *HSF1* depletion reduced global protein translation rate in MPNST cells (Figures 3F, S3E, and S3F), as measured by puromycin labeling.^33,37^ This regulatory relationship was conserved across diverse human cancer cell lines (Figure 3G).

Importantly, treatment with JNK-IN-8, an irreversible JNK inhibitor,^38^ largely restored protein translation following *HSF1* depletion and increased basal translation rates (Figures 3F, S3E, and S3F). Consistent with these observations, in human cancers higher *HSF1* expression correlates with lower JNK activation (Figure S3G), which in turn is associated with elevated mTORC1 activity, as indicated by increased 4EBP1 Ser65 phosphorylation (Figure S3H). In aggregate, our findings demonstrate that JNK activation mediates mTORC1 inhibition and translational suppression in response to *HSF1* deficiency. Thus, by restraining JNK signaling, HSF1 permits robust protein translation fundamental to malignant growth (Figure 3H).

### Lowering protein quantity is an adaptive response to alleviate proteomic instability

We next asked why cells attenuate protein translation in response to HSF1 deficiency. While HSF1 maintains protein quality through HSPs, mTORC1 regulates protein quantity by controlling translation. Because it is beneficial to lower protein quantity when protein quality is compromised, we reasoned that this regulation represents a cellular adaptive response to the imbalance between protein quantity and quality induced by HSF1 deficiency.

Proteomic imbalance disrupts proteome homeostasis and promotes proteomic instability, which may ultimately trigger amyloidogenesis. To induce proteomic imbalance, we first stimulated protein translation in MPNST cells by inhibiting JNK. As expected, treatment with JNK-IN-8 effectively reduced phosphorylation of c-JUN, a canonical JNK substrate,^34^ in MPNST cells (Figure 4A). Consistent with enhanced protein translation, JNK-IN-8 further activated mTORC1 signaling in these cells (Figures 4A and S4A). In contrast, only minimal effects were observed in HSCs (Figure 4B), in line with their low basal JNK activity, as evidenced by unchanged c-JUN phosphorylation. Intriguingly, the heightened protein translation following JNK inhibition was accompanied by elevated amyloid accumulation in MPNST cells (Figures 4C and S4B), whereas no significant changes were observed in HSCs (Figure 4D).

**Figure 4:**
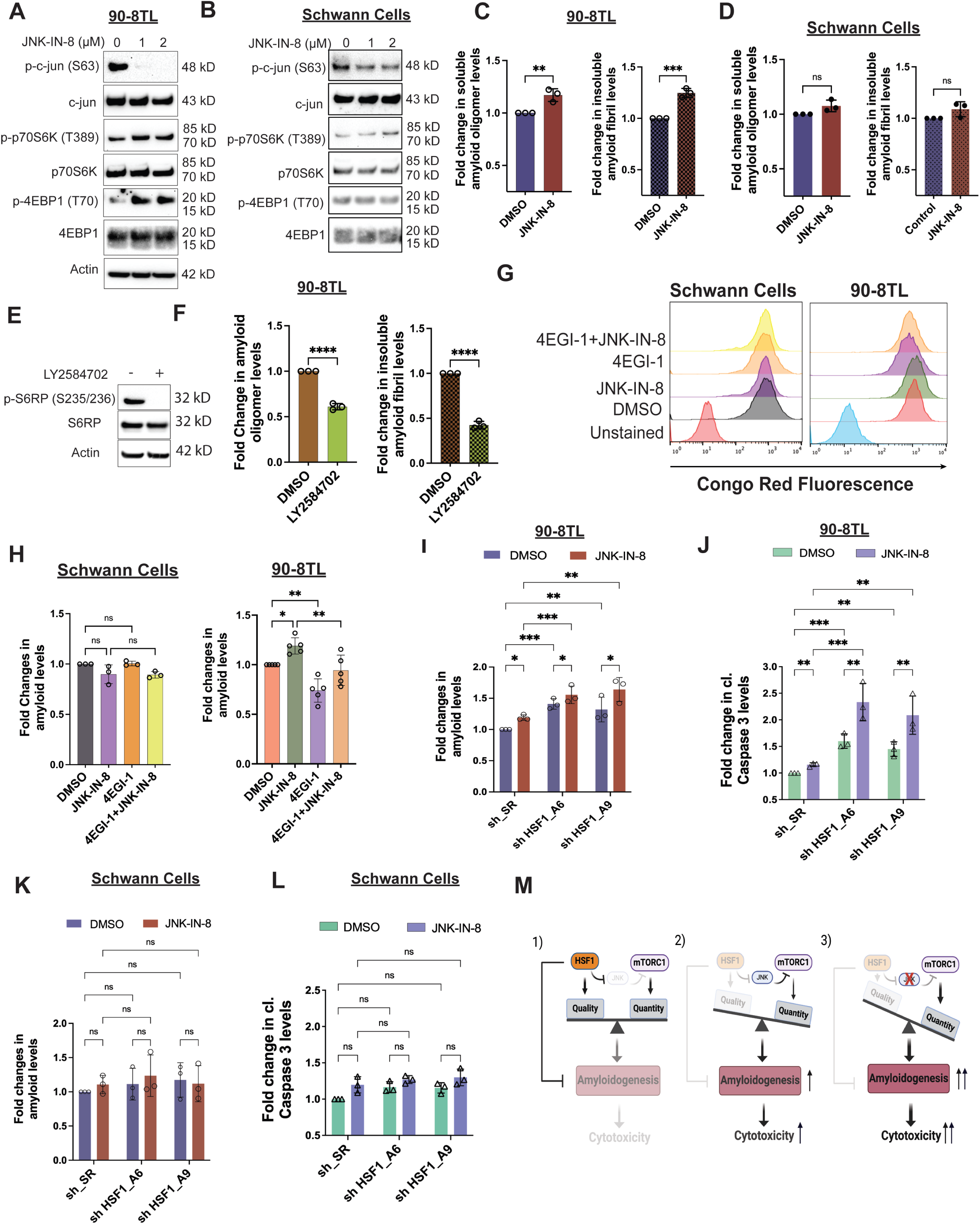
Protein translation is causally related to amyloidogenesis in MPNST cells. (A) and (B) Detection of mTORC1 stimulation following JNK inhibition in 90-8TL and immortalized human Schwann cells by immunoblotting. Cells were treated with JNK-IN-8 for 2 days. (C) and (D) Quantitation of soluble AOs and insoluble AFs by ELISA in 90-8TL and human Schwann cells with and without 3 μM JNK-IN-8 treatment for 3 days (mean ± SD, n=3 independent experiments, Student’s t test). (E) Detection of LY2584702-mediated S6K inhibition in 90-8TL cells by immunoblotting. Cells were treated with 20 μM LY2584702 for 2 days. (F) Quantitation of soluble AOs and insoluble AFs by ELISA in 90-8TL cells with and without 20 μM LY2584702 treatment for 2 days (mean ± SD, n=3 independent experiments, Student’s t test). (G) CR staining of 90-8TL and immortalized human Schwann cells with and without JNK-IN-8 and 4EGI-1 treatment, detected by flow cytometry. (H) Quantitation of (G) using the median fluorescence intensity (mean ± SD, n=3 independent experiments, One-way ANOVA). (I) Quantitation of amyloids by CR staining in 90-8TL cells with and without *HSF1* depletion and 3 μM JNK-IN-8 treatment (mean ± SD, n=3 independent experiments, Two-way ANOVA). (J) Quantitation of caspase 3 cleavage by ELISA in 90-8TL cells with and without *HSF1* depletion and 3 μM JNK-IN-8 treatment (mean ± SD, n=3 independent experiments, Two-way ANOVA). (K) Quantitation of amyloids by CR staining in immortalized human Schwann cells with and without *HSF1* depletion and 3 μM JNK-IN-8 treatment (mean ± SD, n=3 independent experiments, Two-way ANOVA). (L) Quantitation of caspase 3 cleavage by ELISA in immortalized human Schwann cells with and without *HSF1* depletion and 3 μM JNK-IN-8 treatment (mean ± SD, n=3 independent experiments, Two-way ANOVA). (M) Schematic depiction of the adaptation of MPNST cells to *HSF1* depletion by activating JNK to suppress mTORC1 and protein translation, thereby alleviating amyloidogenesis and cytotoxicity.

To directly assess the role of protein translation in amyloidogenesis, we inhibited translation using two pharmacological agents with distinct mechanisms of action: 4EGI-1, which disrupts eIF4E–eIF4G interactions,^39^ and LY2584702, an inhibitor of p70S6K.^40^ Both compounds reduced basal amyloid levels in MPNST cells (Figures 4E,4F, and S4C), establishing a causal link between protein translation and amyloidogenesis. Furthermore, translation inhibition abrogated the induction of amyloids by JNK-IN-8 in MPNST cells (Figures 4G, 4H, S4D, and S4E), indicating that JNK suppresses amyloidogenesis by inhibiting translation. No significant effects were observed in HSCs (Figures 4G and 4H).

Given this relationship, we reasoned that preventing translation attenuation would aggravate proteomic imbalance and enhance cytotoxicity in malignant cells. Consistent with this notion, JNK inhibition by JNK-IN-8 further elevated amyloid levels in *HSF1*-depleted MPNST cells, accompanied by exacerbated apoptosis as evidenced by caspase 3 cleavage (Figures 4I, 4J, and S4F).^41^ Notably, CR treatment suppressed this apoptosis, as indicated by reduced PARP cleavage (Figure S4G),^42^ causally implicating amyloidogenesis in this cell death. In contrast, *HSF1* depletion and JNK inhibition had minimal effects on non-transformed Schwann cells (Figures 4K and 4L), consistent with their balanced proteome and low propensity for amyloid formation. Together, these findings demonstrate that proteomic imbalance in malignant cells drives proteomic instability and amyloidogenesis, leading to cytotoxicity. In non-transformed cells, balanced protein quantity and quality prevent such instability. Thus, attenuation of protein translation represents an adaptive mechanism by which cancer cells mitigate proteotoxic stress and survive proteomic imbalance (Figure 4M).

### mTORC1 stimulation aggravates proteomic imbalance and cytotoxicity induced by HSF1 inhibition

Because JNK regulates multiple targets beyond mTORC1, we sought to stimulate mTORC1 and protein translation specifically. To this end, we genetically inhibited the tumor suppressor Tuberous Sclerosis Complex subunit 2 (*TSC2*), a key negative regulator of mTORC1,^43^ in MPNST cells. As expected, *TSC2* knockdown in MPNST cells increased phosphorylation of both p70S6K and 4EBP1, indicative of mTORC1 activation, which was accompanied by elevated amyloid levels (Figures S5A-S5F). Like genetic *HSF1* depletion, pharmacological inhibition of HSF1 with DTHIB— a small-molecule inhibitor that binds the HSF1 DNA-binding domain to block transcription^44^ —elevated global protein polyubiquitination and amyloids in MPNST cells (Figures 5A, 5B, and S5G), indicating proteomic instability. Notably, HSF1 inhibition induced considerably greater amyloid accumulation in *TSC2*-deficient cells than in control cells (Figure 5B), suggesting that elevated protein translation exacerbates proteotoxic stress. *TSC2*-deficient cells also exhibited heightened JNK activation and mTORC1 inhibition following DTHIB treatment, reflecting an amplified cellular response to severe proteotoxic stress. Correspondingly, whereas control MPNST cells showed only modest cytotoxicity upon HSF1 inhibition, measured by plasma membrane permeabilization, *TSC2*-deficient cells were markedly more sensitive (Figures 5C, 5D, and S5H-S5K). Importantly, either inhibition of protein translation or blockade of amyloid formation with CR largely rescued this cytotoxicity (Figures 5C and 5D), causally implicating proteomic instability and amyloidogenesis in this cell death. In support of a role for HSF1 in enabling *TSC2* deficiency, in tumors harboring *TSC2* deletion higher *HSF1* gene expression correlates with poorer patient survival (Figure S5L).

**Figure 5:**
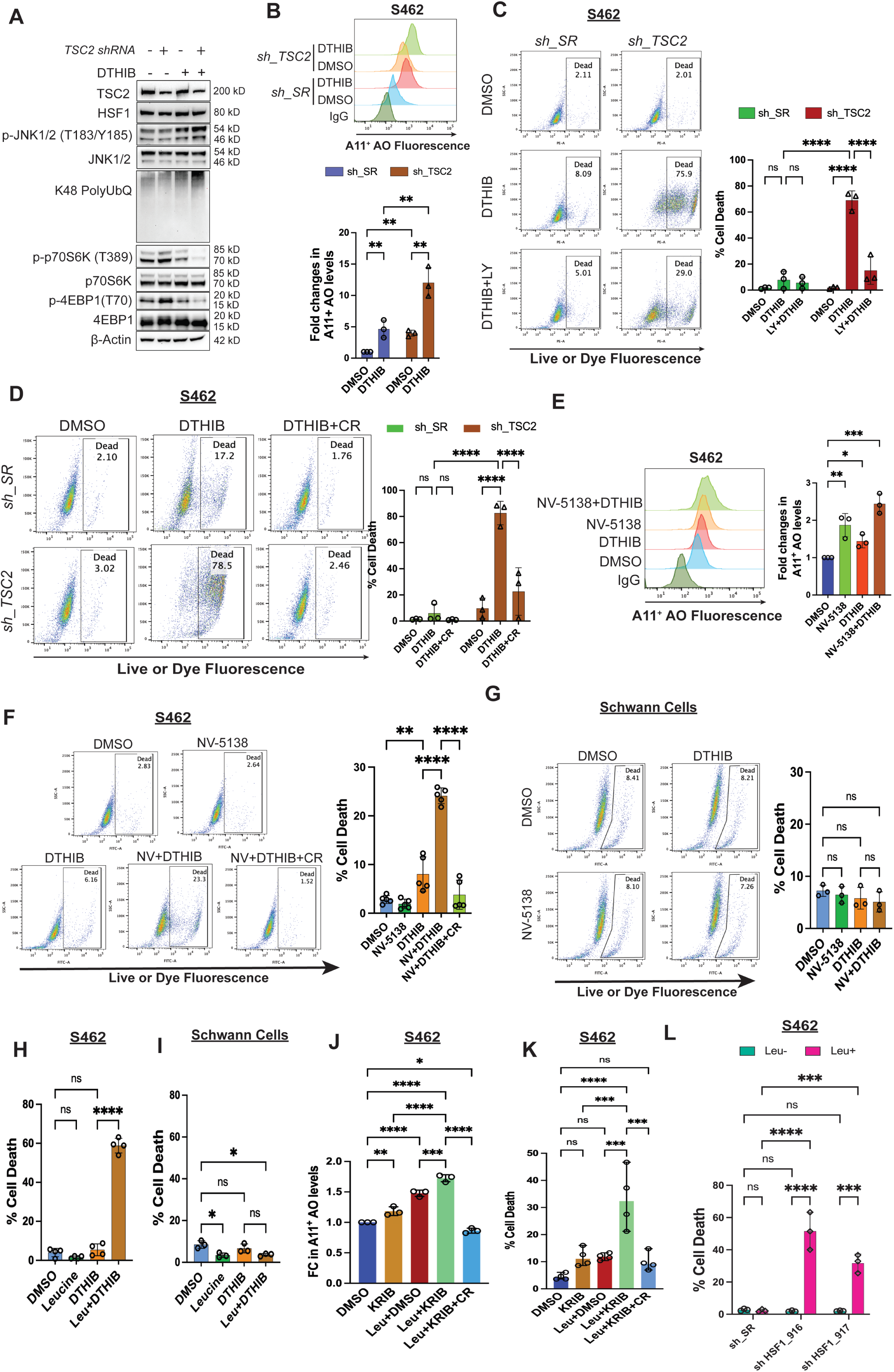
mTORC1 stimulation aggravates the amyloidogenesis and cytotoxicity elicited by HSF1 inhibition. (A) Immunoblotting detection of JNK activation, mTORC1 activity, and protein polyubiquitination in S462 cells following stable *TSC2* knockdown and/or HSF1 inhibition by 10 μM DTHIB for 3 days. (B) Quantitation of soluble AOs by flow cytometry using A11 antibody staining in S462 cells with and without *TSC2* knockdown and 10 μM DTHIB treatment (mean ± SD, n=3 independent experiments, Two-way ANOVA). (C) Quantitation of cytotoxicity by flow cytometry using Live-or-Dye stains in S462 cells with and without stable *TSC2* knockdown and DTHIB (10 μM) or combined DTHIB and LY2584702 (20 μM) treatment (mean ± SD, n=3 independent experiments, Two-way ANOVA). Cells were pre-treated with LY2584702 for 1 day followed by DTHIB treatment for another 3 days. (D) Quantitation of cytotoxicity by flow cytometry using Live-or-Dye stains in S462 cells with and without stable *TSC2* knockdown and DTHIB (10 μM) or combined DTHIB and CR (30 μM) treatment (mean ± SD, n=3 independent experiments, Two-way ANOVA). Cells were pre-treated with CR for 1 day followed by DTHIB treatment for another 3 days. (E) Quantitation of soluble AOs by flow cytometry using A11 antibody staining in S462 cells with and without 500 μM NV-5138 stimulation and 10 μM DTHIB treatment (mean ± SD, n=3 independent experiments, One-way ANOVA). (F) Quantitation of cytotoxicity by flow cytometry using Live-or-Dye stains in S462 cells with and without 500 μM NV-5138 stimulation and 10 μM DTHIB or combined DTHIB and CR treatment (mean ± SD, n=5 independent experiments, One-way ANOVA). (G) Quantitation of cytotoxicity by flow cytometry using Live-or-Dye stains in immortalized human Schwann cells with and without 500 μM NV-5138 stimulation and 10 μM DTHIB treatment (mean ± SD, n=3 independent experiments, One-way ANOVA). (H) Quantitation of cytotoxicity by flow cytometry using Live-or-Dye stains in S462 cells with and without 500 μM L-leucine stimulation and 10 μM DTHIB treatment (mean ± SD, n=4 independent experiments, One-way ANOVA). (I) Quantitation of cytotoxicity by flow cytometry using Live-or-Dye stains in immortalized human Schwann cells with and without 500 μM L-leucine stimulation and 10 μM DTHIB treatment (mean ± SD, n=3 independent experiments, One-way ANOVA). (J) Quantitation of soluble AOs by ELISA in S462 cells treated with and without 800 μM L-leucine and 20 μM KRIBB11 or 30 μM CR (mean ± SD, n=3 independent experiments, One-way ANOVA). (K) Quantitation of cytotoxicity by flow cytometry using Live-or-Dye stains in S462 cells treated as above (mean ± SD, n=4 independent experiments, One-way ANOVA). (L) Quantitation of cytotoxicity by flow cytometry using Live-or-Dye stains in S462 cells stably expressing inducible shRNAs with and without L-leucine stimulation. Cells were first cultured with or without 1.2 mM L-leucine for 2 days, followed by addition of 20 ng/ml doxycycline for 7 days (mean ± SD, n=3 independent experiments, One-way ANOVA).

To exclude potential mTORC1-independent effects of *TSC2* deficiency, we stimulated mTORC1 using NV-5138, a leucine analog currently in clinical trials as an antidepressant.^45,46^ Amino acids, particularly leucine, are potent activators of mTORC1.^47^ Of note, although NV-5138 activates mTORC1, it cannot be incorporated into protein synthesis.^45,46^ To maximize mTORC1 activation *ex vivo*, cells were cultured in leucine-free medium. Similar to *TSC2* knockdown, NV-5138 stimulated mTORC1 and elevated amyloid levels in MPNST cells (Figures S5M and 5E). However, it elicited pronounced cytotoxicity only when combined with HSF1 inhibition, which was largely rescued by CR treatment (Figure 5F). In contrast, combined NV-5138 and DTHIB treatment had minimal effects on non-transformed Schwann cells (Figure 5G), consistent with their reduced responsiveness to mTORC1 stimulation (Figures S5N). We next reasoned that by directly contributing to protein quantity, L-leucine might impact the cancer proteome more profoundly. In both S462 and 90-8TL cells, L-leucine synergized with DTHIB to induce severe cytotoxicity (Figures 5H and S5O), whereas no cytotoxicity was induced in HSCs (Figure 5I).

To further corroborate the HSF1-dependent effects of DTHIB, we tested an independent small-molecule inhibitor, KRIBB11^48^, which binds the HSF1 transactivation domain to block the recruitment of pTEFb. KRIBB11, consistent with DTHIB, synergized with L-leucine to promote amyloid formation and trigger severe cytotoxicity in MPNST cells; importantly, CR treatment both diminished amyloid levels and blocked cell death (Figures 5J and 5K). Finally, genetic depletion of *HSF1* using shRNAs similarly synergized with L-leucine stimulation to provoke prominent cytotoxicity (Figure 5L), providing orthogonal validation.

Collectively, these findings demonstrate that mTORC1 stimulation synergizes with HSF1 inhibition to drive severe cytotoxicity in cancer cells. This effect is mediated, at least in part, by unconstrained proteomic imbalance and amyloidogenesis.

### Unconstrained proteomic imbalance incites pronounced cell death

HSF1 inhibition alone induced both apoptotic and non-apoptotic death in MPNST cells, as indicated by the presence or absence of caspase 3 cleavage. Intriguingly, concurrent mTORC1 stimulation shifted cell death predominantly toward a non-apoptotic mode, characterized by plasma membrane permeabilization without caspase 3 cleavage (Figures 6A and 6B).

**Figure 6:**
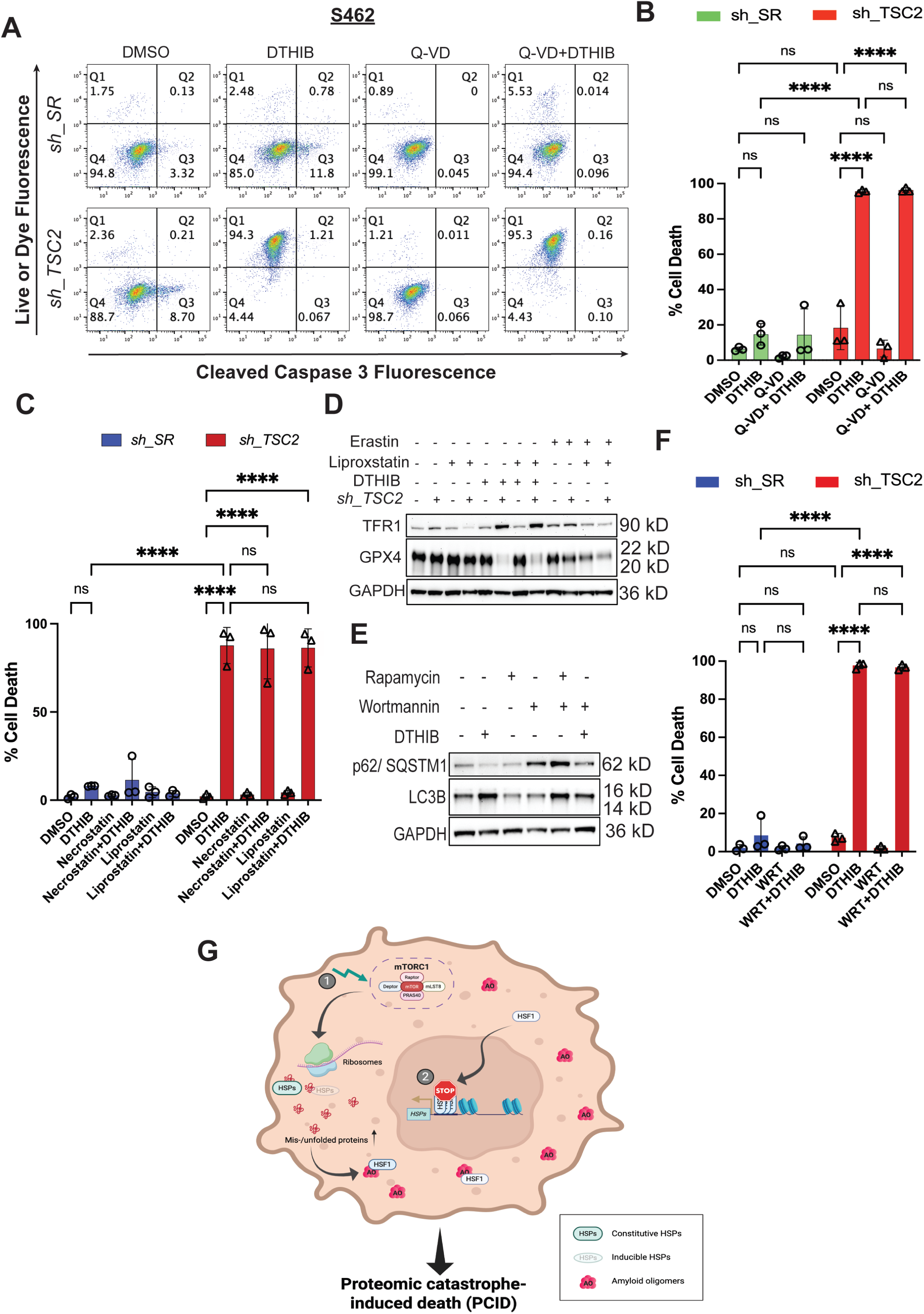
Concurrent HSF1 inhibition and mTORC1 stimulation instigate widespread non-apoptotic cell death. (A) and (B) Quantitation of cytotoxicity by flow cytometry using Live-or-Dye stains combined with cleaved caspase 3 (Asp175) antibody staining. S462 cells with and without stable *TSC2* knockdown were treated with DMSO, 10 μM DTHIB, or combined DTHIB and 30 μM Q-VD-OPh (mean ± SD, n=3 independent experiments, Two-way ANOVA). (C) Quantitation of cytotoxicity by flow cytometry using Live-or-Dye stains in S462 cells with and without stable *TSC2* knockdown. Cells were treated with 10 μM DTHIB alone or co-treated with 10 μM DTHIB and 30 μM Necrostatin-1 or 20 μM Liproxstatin-1 for 3 days (mean ± SD, n=3 independent experiments, Two-way ANOVA). (D) Immunoblotting detection of ferroptosis markers in S462 cells with and without stable *TSC2* knockdown treated with 10 μM DTHIB alone or co-treated with 10 μM DTHIB and 20 μM liproxstatin-1 for 3 days. Erastin was included as a positive control to induce canonical ferroptosis. (E) Immunoblotting detection of autophagy markers in S462 cells with stable *TSC2* knockdown treated with 10 μM DTHIB alone or co-treated with 10 μM DTHIB and 3 μM wortmannin for 3 days. Rapamycin was included as a positive control to induce autophagy. (F) Quantitation of cytotoxicity by flow cytometry using Live-or-Dye stains in S462 cells with and without stable *TSC2* knockdown. Cells were treated with 10 μM DTHIB alone or co-treated with 10 μM DTHIB and 3 μM wortmannin for 3 days (mean ± SD, n=3 independent experiments, Two-way ANOVA). (G) Schematic depiction of instigation of cell death by severe proteomic imbalance, owing to simultaneous mTORC1 stimulation (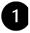) and HSF1 inhibition (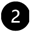). In cancer cells, constitutive HSF1 activation provides extra chaperoning capacity to cope with elevated protein misfolding, partly due to enhanced protein synthesis and widespread genetic mutations. Nevertheless, amyloids still emerge, although at low levels. Importantly, HSF1 can neutralize highly toxic amyloid oligomers, averting lethal consequences. By contrast, HSF1 inhibition diminishes chaperoning capacity, insufficient to counterbalance the robust protein translation. mTORC1 stimulation further aggravates this proteomic imbalance, which, in turn, promotes amyloidogenesis and proteomic catastrophe. In consequence, the amounts of amyloid oligomers surpass the neutralizing capacity of HSF1, leading to cell death; nonetheless, it remains unclear how this proteomic catastrophe-induced death (PCID) occurs.

Multiple forms of non-apoptotic cell death, including pyroptosis, necroptosis, ferroptosis, and autophagic cell death, can lead to loss of membrane integrity. To define the mechanism, we first tested the role of caspases. Corroborating a caspase-independent mechanism, the pan-caspase inhibitor Q-VD-OPh failed to prevent membrane permeabilization in *TSC2*-deficient MPNST cells treated with the HSF1 inhibitor DTHIB (Figures 6A and 6B).^49^ In contrast, Q-VD-OPh effectively blocked apoptosis in control MPNST cells (Figures 6A and S6A), confirming its activity. Because Q-VD-OPh also inhibits caspase 1 and no caspase 1 cleavage was detected (Figure S6B),^50^ a key mediator of pyroptosis,^51^ these results argue against pyroptotic cell death. Similarly, HSF1 inhibition did not induce necroptosis in *TSC2*-deficient MPNST cells, as indicated by the absence of MLKL Ser358 phosphorylation (Figure S6B), a hallmark of necroptotic execution.^52^ Congruently, the necroptosis inhibitor necrostatin-1 had no effects on membrane permeabilization (Figures 6C and S6C).^52^

Intriguingly, HSF1 inhibition in *TSC2*-deficient MPNST cells decreased glutathione peroxidase 4 (GPX4) but increased transferrin receptor 1 (TFR1), accompanied by markedly elevated lipid peroxidation (Figures 6D and S6D), features typically associated with ferroptosis.^53,54^ However, the ferroptosis inhibitor liproxstatin-1,^53,54^ while effectively reversing TFR1 elevation induced by the ferroptosis inducer erastin, failed to suppress either TFR1 upregulation or membrane permeabilization triggered by DTHIB (Figures 6C, 6D, and S6C). These results are not due to ineffective inhibition, as both necrostatin-1 and liproxstatin-1 robustly blocked necroptosis and ferroptosis, respectively, in control HT-29 cells (Figure S6E). HSF1 inhibition also elicited autophagy, as evidenced by diminished p62/SQSTM1 and increased LC3B-II levels (Figure 6E).^55^ To assess whether autophagy contributes to cell death, we blocked autophagy using the class III PI3K inhibitor wortmannin.^56^ Wortmannin effectively blocked autophagic flux, as shown by reversal of rapamycin-induced changes in p62/SQSTM1 and LC3B-II (Figure 6E), confirming successful inhibition of autophagy. Nevertheless, it failed to prevent membrane permeabilization in *TSC2*-deficient cells treated with HSF1 inhibitors (Figures 6F and S6F).

These findings indicate that ferroptosis- and autophagy-associated changes occur downstream of severe proteotoxic stress and are not the primary causes of cell death. Instead, our data support a model in which unrestrained proteomic instability and consequent amyloidogenesis trigger a unique form of cell death that is independent of apoptosis, autophagy, and canonical programmed necrosis pathways (Figure 6G). We term this process <u>p</u>roteomic <u>c</u>atastrophe-induced <u>d</u>eath (PCID). While its precise molecular execution remains to be fully defined, PCID represents a mechanistically distinct vulnerability arising from the collapse of proteome homeostasis in cancer cells.

### Unconstrained proteomic imbalance suppresses *in vivo* tumor growth

Due to the limited *in vivo* bioavailability of DTHIB in our mouse models, we employed KRIBB11 for *in vivo* studies. In agreement with our *ex vivo* findings, combined L-leucine supplementation and KRIBB11 administration significantly suppressed the growth of xenografted human S462 MPNST tumors in immunocompromised mice (Figure 7A). Despite this overall reduction in tumor burden, 2 out of 8 tumors did not respond to the combination treatment (Figure S7A), resulting in notable variations and a right-skewed distribution. Notably, KRIBB11 alone only marginally affected tumor growth; conversely, in the presence of leucine it markedly slowed tumor exponential growth rates and delayed tumor volume doubling time (Figures S7B). This combination treatment also trended toward improved survival (Figure S7C).

**Figure 7:**
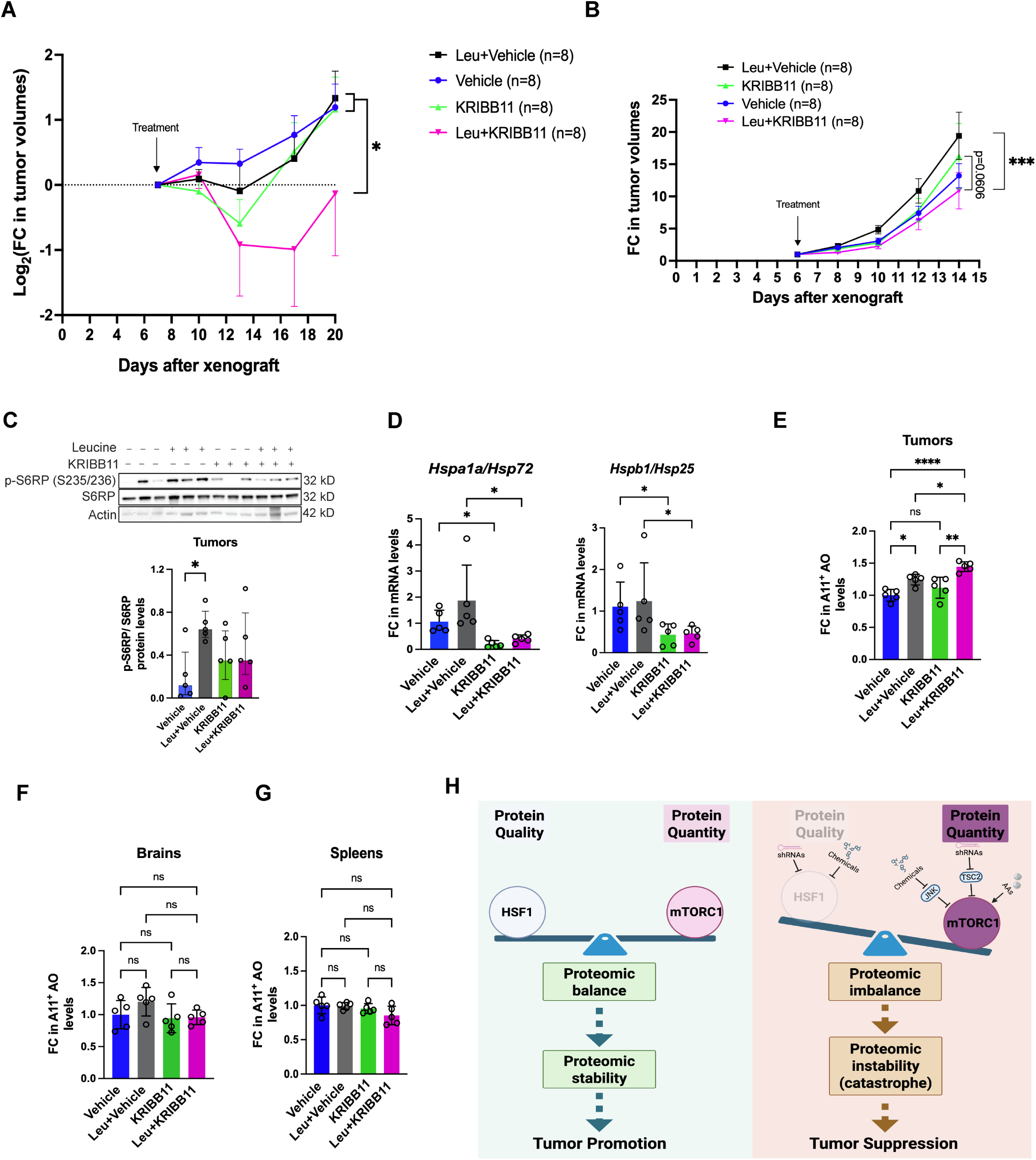
Concurrent HSF1 inhibition and mTORC1 stimulation suppress *in vivo* tumor growth. (A) Combined KRIBB11 treatment and L-leucine stimulation impaired the *in vivo* growth of xenografted S462 human MPNST cells in NIH-III nude mice (mean ± SEM, n=8 mice per group, log_2_ transformation for Two-way ANOVA). (B) Combined KRIBB11 treatment and L-leucine stimulation impeded the *in vivo* growth of xenografted B16-F10 murine melanoma cells in C57BL/6J mice (mean ± SEM, n=8 mice per group, log_2_ transformation for Two-way ANOVA). (C) Quantitation of phospho-S6RP by immunoblotting in treated melanomas. Images show representative 3 tumors each group (mean ± SEM, n=5 tumors per group, One-way ANOVA). (D) Quantitation of *Hsp72* and *Hsp25* mRNAs in treated melanomas by RT-qPCR (mean ± SD, n=5 tumors per group, One-way ANOVA). (E) Quantitation of soluble AOs in treated B16F10 melanomas by ELISA ((mean ± SD, n=5 tumors per group, One-way ANOVA). (F) and (G) Quantitation of soluble AOs in brains and spleens from treated C57BL/6J mice by ELISA (mean ± SD, n=5 brains or spleens per group, One-way ANOVA). (H) Schematic depiction of the concept of provoking proteomic catastrophe to combat malignancy. On the one hand, in cancer cells, mTORC1 is inevitably activated to stimulate protein translation, markedly augmenting protein quantity. On the other hand, the extra chaperoning capacity governed by HSF1, albeit dispensable for normal life, becomes necessary to ensure sufficient protein quality in cancer cells, thereby counterbalancing augmented protein quantity and suppressing proteomic instability. Thus, proteomic balance promotes malignant growth. By contrast, disrupting proteomic balance, through HSF1 inhibition, is sufficient to elicit proteomic instability and tumor suppression. However, simultaneous mTORC1 stimulation can remarkably drive proteomic imbalance, causing proteomic catastrophe that leads to tumor suppression.

To evaluate the broader applicability of this therapeutic strategy, we established a syngeneic mouse melanoma model using B16-F10 cells in C57BL/6J mice. Similar to the MPNST model, combined KRIBB11 and L-leucine supplementation significantly impeded tumor growth in this aggressive model (Figures 7B and S7D). As expected, L-leucine activated mTORC1 signaling, as indicated by increased S6RP phosphorylation (Figure 7C). Successful inhibition of HSF1 by KRIBB11 was confirmed by the reduced transcription of *Hspa1a/Hsp72* and *Hspb1/Hsp25* within the tumors (Figure 7D). Significantly, the combination treatment markedly induced amyloid accumulation within tumors (Figure 7E). In stark contrast, no amyloids were induced in normal tissues, including the brain and spleen (Figures 7F and 7G). Moreover, these treatments did not cause significant loss of body weight (Figure S7E). In aggregate, these *in vivo* findings from two distinct mouse models demonstrate that enforced proteomic imbalance—achieved through combined HSF1 inhibition and mTORC1 stimulation—induces proteomic catastrophe that suppresses tumor growth while sparing normal tissues (Figure 7H).

## DISCUSSIONS

Genomic instability is widely recognized as a fundamental driver of oncogenesis. In contrast, the role of proteomic instability in cancer remains obscure.

### Proteomic instability is tumor suppressive yet inevitable

Proteomic instability does occur in cancer, as evidenced by elevated protein polyubiquitination, aggregation, and even amyloidogenesis. Multiple factors contribute to this instability, including protein oxidation by reactive oxygen species, mutation-induced protein misfolding, and increased protein dosage resulting from aneuploidy.^1,3^ Our recent findings further identify uncontrolled protein translation as a key contributor to amyloidogenesis. This is not entirely surprising, as it is estimated that up to 30% of newly synthesized polypeptides are misfolded and targeted for proteasomal degradation.^57^ Aberrant protein synthesis is therefore likely to increase the burden of misfolded proteins, promoting protein aggregation and amyloid formation—particularly in cancer cells, which harobor numerous mutations affecting protein folding. Heightened protein translation, paradoxically, is also essential for malignant growth, rendering proteomic instability and amyloidogenesis an unavoidable consequence of oncogenesis.

At the cellular level, amyloids are toxic to both neurons and cancer cells. At the organismal level, while amyloidogenesis contributes to neurodegeneration, it appears to exert tumor-suppressive effects. Mechanistically, our findings demonstrate that soluble AOs can directly target HSP60, a key guardian of the mitochondrial proteome. It is likely that additional targets and mechanisms contribute to amyloid toxicity in cancer. Moreover, the composition of cancer-associated amyloids remains largely undefined. Beyond well-characterized species such as Alzheimer’s disease-associated Aβ, identifying the proteins that form amyloids in cancer will be critical for understanding their functional roles and therapeutic potential.

### HSF1 contains proteomic instability to enable malignancy

The diverse pro-oncogenic functions of HSF1 have been attributed to a wide array of mechanisms; nonetheless, a unifying principle has remained elusive.

As an inherent consequence of malignant transformation, proteomic instability must be restrained to avert tumor suppression and sustain cancer cell fitness. Now, our findings identify containment of proteomic instability—particularly amyloidogenesis—as a central, unifying pro-oncogenic function of HSF1. Notably, HSF1 does not eliminate amyloid formation but instead limits its extent and toxicity. This is consistent with the fact that uncontrolled protein translation, a hallmark of cancer, continuously drives amyloidogenesis. Consequently, cancer cells become dependent on HSF1 to buffer this intrinsic proteotoxic burden, providing a mechanistic basis for “HSF1 addiction”.

Mechanistically, HSF1 contains amyloidogenesis through both indirect and direct means. Indirectly, HSF1 induces HSPs to ensures proper protein folding and maintain proteome integrity, thereby limiting the generation of misfolded proteins and amyloids. This is supported by the observation that HSF1 inhibition leads to increased protein ubiquitination, aggregation, and amyloid accumulation. Directly, HSF1 neutralizes soluble AOs once they form. With HSF1 inhibition, AOs are free to target HSP60, compromising the mitochondrial proteome and triggering cytoxicity. Thus, in cancer cells HSF1 inhibition not only increases the burden of amyloids but, critically, exposes their intrinsic toxicity. Together, these findings establish that HSF1 enables malignancy by buffering proteomic instability, positioning it as a central safeguard against proteotoxic collapse in cancer cells.

### mTORC1 signaling becomes tumor suppressive in the absence of HSF1

Given the fundamental dependency of malignant growth on protein translation, mTORC1 signaling is frequently hyperactivated in human cancers and is traditionally viewed as oncogenic.^58,59^ However, our findings reveal a striking context-dependent shift: in the absence of HSF1, mTORC1 signaling becomes tumor suppressive. This functional reversal arises from a profound imbalance between protein quantity and quality. Without HSF1 to maintain proteome integrity, continued mTORC1-driven protein synthesis exacerbates proteomic instability, leading to enhanced amyloidogenesis and cytotoxicity. The observation that cytotoxicity is alleviated by either translation inhibitors or blockade of amyloid formation with CR strongly supports this model.

Conceptually, HSF1 enables the oncogenic potential of mTORC1 signaling by buffering the proteotoxic stress associated with elevated protein synthesis. In its absence, mTORC1-driven translation becomes detrimental, converting a canonical growth-promoting pathway into a tumor-suppressive force. These findings further establish HSF1 as a critical oncogenic enabler that permits cancer cells to tolerate the proteomic burden of malignant growth.

### Inducing proteomic catastrophe represents a next-generation therapeutic paradigm

Conventional chemo- and radiotherapies primarily target the cancer genome by inducing DNA damage. An emerging paradigm in oncology is to instead target the cancer proteome. To date, most efforts have focused on inhibiting individual HSP families, such as HSP70 and HSP90.^60,61^ However, this strategy faces intrinsic limitations: cells harbor both constitutive HSPs, which maintain essential basal proteostasis, and inducible HSPs, which provide additional chaperoning capacity that is dispensable for basal proteostasis but required under stress. The structural and functional similarity between these HSPs complicates selective targeting, and redundancy across multiple HSP families further limits the efficacy of targeting any single chaperone system.

Our findings, together with prior studies, suggest that malignant cells operate under chronic proteotoxic stress, resulting in constitutive HSF1 activation. Consequently, cancer cells become reliant on the inducible chaperoning capacity governed by HSF1 for growth and survival. Targeting HSF1 therefore represents a strategy to selectively disable this stress-adaptive reserve while preserving basal proteostasis in normal cells. These insights provide a strong rationale for therapeutic targeting of HSF1, an approach that is now beginning to enter clinical investigation.^62^

In response to HSF1 inhibition, cancer cells attenuate protein synthesis as an adaptive mechanism, highlighting the importance of maintaining a balance between protein quantity and quality. This observation led us to explore whether deliberately driving proteomic imbalance could be therapeutically exploited.

Although mTORC1 hyperactivation is widespread in human cancers and has long been targeted with inhibitors,^59, 63^ these agents have largely produced cytostatic effects and limited clinical benefit.^63,64^ Moreover, because protein synthesis is essential for normal cellular function, systemic mTORC1 inhibition carries significant toxicity, including immunosuppression.^63,65^

In contrast to conventional approaches, we pursued a counterintuitive strategy: stimulating mTORC1 to drive proteomic imbalance. When combined with HSF1 inhibition, mTORC1 stimulation engenders catastrophic proteomic instability in cancer, culminating in cell death and tumor suppression (Figure 7G). This PCID offers a wider therapeutic window, as normal cells, with intact proteostasis and low proteotoxic stress, are largely spared. Mechanistically, mTORC1 activation increases protein synthesis, which can be accommodated in normal cells but overwhelms the already stressed proteome of cancer cells, particularly given their burden of mutant and misfolded proteins. Although cancer cells attempt to adapt via JNK-mediated suppression of mTORC1, further specific blockade of this adaptive response is expected to amplify proteomic imbalance and enhance therapeutic efficacy.

Transcription factors, traditionally, have been considered “undruggable” targets for small molecules. While prototype direct HSF1 inhibitors such as DTHIB and KRIBB11 provide important proof-of-concept, their limited *in vivo* potency underscores the need for more clinically viable compounds. Nonetheless, our study establishes a new therapeutic paradigm: the deliberate induction of proteomic catastrophe in cancer cells. We demonstrate a feasible strategy to achieve this by simultaneously disrupting proteome quality control and enhancing protein synthesis, thereby overwhelming the proteostatic capacity of malignant cells. This approach may be broadly applicable across tumor types and represents a promising next-generation strategy for cancer therapy.

## ACKNOWLEDGMENTS

We would like to thank the Optical Microscopy and Image Analysis lab (OMAL) for their assistance with the confocal microscopy studies and Holly Morris for her assistance in animal studies. Schematic illustrations are created with BioRender.com. This work was supported by the grant from Department of Defense Congressionally Directed Medical Research Programs to C.D. (NF150028) and by the Intramural Research Program of the NIH, National Cancer Institute, Center for Cancer Research (1ZIABC011767). The content of this publication does not necessarily reflect the views or policies of the Department of Health and Human Services, nor does mention of trade names, commercial products, or organizations imply endorsement by the U.S. government.

## AUTHOR CONTRIBUTIONS

B.M.R., O. S., and K-H.C. designed and conducted the experiments. C.D. conceptualized and supervised this study and analyzed the results. B.M.R. and C.D. wrote the manuscript.

## EXPERIMENTAL MATERIALS AND METHODS

### Cells, Chemicals, and reagents

S462 (female) and 90-8TL (female) cells were kindly provided by Dr. Karen Cichowski. sNF96.2 cells (male) and immortalized human Schwann cells (hTERT ipn02.3 2λ, female) were purchased from ATCC. Primary human Schwann cells (Cat#1700, ScienCell Research Laboratories) were grown on poly-L-lysine coated T-75 flasks and maintained in complete Schwann Cell Medium (Cat#1701, ScienCell Research Laboratories). B16-F10-luc2 (male) cells were purchased from ATCC (cat#CRL-6475-LUC2). All cell lines, except 90-8TL and HSCs (no known STR profiles available), were authenticated by ATCC. All cell lines, except primary human Schwann cells, were maintained in DMEM supplemented with 10% HyClone bovine growth serum and 1% penicillin–streptomycin. Cells were maintained in an incubator with 5% CO2 at 37 °C.

The following chemicals were purchased from commercial sources: JNK-IN-8 (Cat#S4901, Selleckchem), 6-FAM*-*dC*-*puromycin (cat#NU-925-6FM, Jena Bioscience), Congo red (cat#B2431014, Thermo Scientific Chemicals), Thioflavin T (cat#211760050, Thermo Scientific Chemicals), LY2584702 (cat#S7698, Selleckchem), 4EGI-1 (cat#S7369, Selleckchem), and DTHIB (cat# HY-138280, MedChemExpress), KRIBB11 (cat#HY-100872, MedChemExpress), KRIBB11 (cat#T3652, TargetMol Chemicals), Q-VD-OPh (cat#HY-12305, MedChemExpress), necrostatin-1 (cat#HY-15760, MedChemExpress), Liproxstatin-1 (cat#HY-12726, MedChemExpress), wortmannin (cat#HY-10197, MedChemExpress), rapamycin (cat#HY-10219, MedChemExpress), erastin (cat#HY-15763, MedChemExpress), MHY1485 (cat#HY-B0795, MedChemExpress), NV-5138 HCl (cat#HY-114384B, MedChemExpress), and Liperfluo (cat#L248, Dojindo Laboratories), dimethylacetamide (cat#T19439, TargetMol Chemicals Inc.), PEG300 (cat#T7022, TargetMol Chemicals Inc.), and Captisol (cat#RC-0C7-100, CyDex Pharmaceuticals).

Rabbit phospho-4EBP1 S65/T70 antibody (cat#sc-12884-R), mouse monoclonal HSF1 (E-4, cat#sc-17757) antibody, rabbit HSF1 antibody (H-311, cat#sc-9144), mouse monoclonal JNK (D-2, cat#sc-7345), rabbit JNK1/3 (C-17, cat#sc-474) antibody were purchased from Santa Cruz Biotechnology; antibodies for phosphor-4EBP1 Thr70 (cat#9455), phospho-4EBP1 Thr37/46 (236B4, cat#2855), phospho-p70S6K T389 (108D2, cat# 9234), phosphor-S6RP Ser235/236 (D57.2.2E, cat#4858), p70 S6K (49D7, cat#2708), 4EBP1 (53H11, cat#9644), S6RP (5G10, cat#2217), mTOR (7C10, cat#2983), phospho-JNK1/2 T183/Y185 (81E11, cat#4668), JNK2 (56G8, cat#9258), c-Jun (60A8, cat#9165), phospho-c-Jun Ser63 II (cat#9261), PARP (cat#9542), Tuberin/TSC2 (D93F12, cat#4308), SQSTM1/p62 (D1Q5S, cat#39749), cleaved caspase 3 (Asp175) (5A1E, cat#9664), caspase 1(D7F10, cat#3866), gasdermin D (E9S1X, cat#39754), phosphor-MLKL Ser358 (D6H3V, cat#91689), GPX4 (E5Y8K, cat#59735), and transferrin receptor/CD71 (H68.4, cat#46222) antibody were purchased from Cell Signaling Technologies; mouse β-actin antibody-HRP (Cat#A00730) conjugates were from GenScript. Anti-K48-specific ubiquitin antibody (Apu2, cat#05-1307) and mouse monoclonal anti-RAPTOR antibody (1H6.2, cat#05-1470) were purchased from EMD Millipore. Mouse monoclonal anti-JNK1 (C-terminal region) antibody (cat#JM2671) was purchased from ECM Biosciences. Anti-amyloid oligomers (A11, cat#SPC-506D) and anti-amyloid fibrils (OC, cat#SPC-507D) antibodies were purchased from StressMarq Biosciences Inc. Mouse monoclonal anti-HSP60 antibody (LK-1, cat#ADI-SPA-806) was purchased from Enzo Life Sciences Inc. β-Actin (cat#GTX629630), GAPDH (cat#GTX627408), α-tubulin (GT114, cat#GTX628802), and LC3B antibody (cat#GTX127375) was purchased from GeneTex Inc.

## METHOD DETAILS

### Lentiviral Particle Production and Transduction of lentiviral shRNAs

Lentiviral pLKO.1 shRNA plasmids targeting human *HSF1* were described previously^10^: TRCN0000007480 (hA6) and TRCN0000007483 (hA9). The sequence of control hairpins targeting a scrambled sequence with no known homology to any human genes is 5′-CCTAAGGTTAAGTCGCCCTCG-3′. The pLKO.1 shRNA plasmid targeting human *TSC2* was a gift from Do-Hyung Kim (Cat#15478, Addgene). Lentiviral pLKO shRNA plasmids were transiently co-transfected individually along with plasmids encoding ΔVPR and VSV-G into HEK293T cells using TurboFect™ (Cat#R0531, Thermo Scientific). Viral supernatants were collected after centrifugation at 1,000 rpm for 15 min. Viral transduction was achieved by incubating target cells with viral supernatants containing 10μg/ml polybrene (Cat#TR-1003-G, Millipore-Sigma) for 24h. For stable knockdown of *TSC2,* lentivirus particles were prepared using pLKO shRNA plasmids targeting human *TSC2*. Scrambled shRNA was used for stable control cells. After 4 days of transduction, cells were gradually selected using 1-4 μg/ml puromycin for 2 weeks.

### Soluble and Insoluble Protein Fractionation

Equal numbers of cells were incubated with cold cell-lysis buffer (100 mM NaCl, 30 mM Tris-HCl pH 7.6, 1% Triton X-100, 20 mM sodium fluoride, 1mM EDTA, 1mM sodium orthovanadate, and 1x Halt™ protease inhibitor cocktail) on ice for 20 min. The crude lysates were first centrifuged at 500xg for 5 min at 4°C. The supernatants from low-speed (500xg) centrifugation were transferred to a new Eppendorf tube. The first pellets were resuspended in DNA digestion solution (2 units DNase I, 1% Triton X-100, 10 mM Tris-HCl, 2.5 mM MgCl2, 0.5 mM CaCl2, pH 7.6) for 30 min at RT. An equal volume of 1xPBS with 2% SDS was then added to resuspend the first pellets for additional 30 min at RT. Following the centrifugation at 17,000xg for 10 min at RT, the supernatants were removed. The supernatants from low-speed centrifuge were added back to the second pellets, followed by centrifugation at 17,000xg for 15 min at 4°C. The final supernatants and pellets from high-speed (17,000xg) centrifugation were collected as detergent-soluble and -insoluble fractions, respectively. Insoluble fractions were further sonicated in 1xPBS with 2% SDS at high intensity using a Bioruptor® Sonication Device (Diagenode) for SDS-PAGE.

### ThT and CR Staining for Flow Cytometry

After washing with 1xPBS, cells were fixed with 4% formaldehyde at RT for 10 min. Following fixation and 1xPBS washing, cells were resuspended in 1 ml permeabilization buffer (1xPBS with 0.5% Triton X-100 and 3 mM EDTA) and incubated on ice for 30 min. Following washing with 1xPBS, Cells were stained with 10 μM ThT or CR dissolved in 1xPBS for 5 min. Cells were washed and resuspended in PBS and fluorescence signals were measured by a FACSCalibur, LSRFortessa X-20 or LSR ΙΙ flow cytometer (BD Biosciences).

### Amyloid Oligomer and Fibril Quantitation by Direct ELISA and Flow Cytometry

To quantitate soluble amyloid prefibrillar oligomers, soluble cell lysates diluted in 1xPBS from equal numbers of cells were coated on each well in a 96-well ELISA plate at 4°C overnight followed by blocking (5% non-fat milk in TBS-0.05% Tween® 20) at RT for 1 hr. Each well was incubated with 100 μl amyloid oligomer antibodies (A11, 1:1,000 diluted in the blocking buffer) at RT for 2 hr. After washing with TBS-T, goat anti-rabbit secondary Ab HRP conjugates (1:1,000 diluted in blocking buffer) were added to each well and incubated at RT for 1 hr. Following washing, 100 μl 1-Step Ultra TMB-ELISA substrates (Cat#34029, Thermo Scientific) were added to each well. After adding 100μl Stop solution, absorbance was measured at 450 nm using a SpectraMax iD5 microplate reader (Molecular Devices).

To quantitate amyloid fibrils, detergent-insoluble fractions from equal numbers of cells were extracted as described in Soluble and Insoluble Protein Fractionation. The final pellets were collected as detergent-insoluble fractions and solubilized by sonication for 15 min in 1xPBS with 2% SDS. Solubilized proteins diluted in 1xPBS were added to each well and incubated at 37°C without cover overnight to air dry the wells. The following steps were identical to the oligomer detection except for the use of amyloid fibril antibodies (OC) as the primary Ab.

For amyloid oligomer and fibril quantification by flow cytometry, cells were collected by trypsinization and fixed in 4% Formaldehyde at RT for 10 min. Following permeabilization using 0.3% Triton-X-100, samples were incubated with primary antibodies A11-FITC (1:100 in 5% BSA, StressMarq Biosciences, cat# SPC-506D-FITC) or OC-PerCP (1:100 in 5% BSA, StressMarq Biosciences, cat# SPC-507D-PCP) at RT for 3 hrs. The cells were washed with PBS and analyzed using flow cytometry.

### Measurement of Cell Viability

Cell viability was determined using the CellTiter-Blue® Cell Viability Assay (cat#G8080, Promega). Cells were grown in 96 well plates (1×10^4^ per well) for 24h. A 20μl CellTiter-Blue® reagent was added to 100μl of media in each well. The plates were incubated for 2h at 37°C and the fluorescence was recorded at 560/590 nm. The reagents were discarded, and the wells were washed with PBS before adding fresh media. Similarly, subsequent readings were recorded at 24h intervals till 96h time point.

### Proximity Ligation Assay

Cells were fixed with 4% formaldehyde in PBS for 15 min at RT. After blocking with 5% normal goat serum in PBS with 0.3% Triton X-100, primary antibodies 1:100 diluted in blocking buffer were incubated with fixed cells overnight at 4°C. Following washing with PLA wash Buffer A, samples were incubated with 1:5 diluted Duolink® in Situ PLA Probe anti-rabbit Plus (cat#DUO92002, Sigma-Aldrich) and anti-mouse Minus (cat#DUO92004, Sigma-Aldrich) at 37°C for 1 hr. Subsequently, ligation (30 minutes, 37°C), rolling circle amplification, and detection (120 minutes, 37°C) were performed using the Duolink® In Situ Detection Reagents Red (cat#DUO92008, Sigma-Aldrich). Nuclei were counterstained with Hoechst 33342. PLA signals were documented by a Zeiss LSM780 confocal microscope (Carl Zeiss). For PLA flow cytometry, an equal number of cells were collected in microcentrifuge tubes by trypsinization and fixed with 4% formaldehyde in PBS for 10 min at RT. Subsequent PLA steps were followed as described earlier. The PLA signals were quantified by a FACSCalibur flow cytometer (BD Biosciences).

Mouse anti-JNK1 (JM2671) and rabbit anti-HSF1 (H-311) antibodies were used for JNK-HSF1 PLA. Rabbit anti-JNK1/3 (C-17) and mouse anti-RAPTOR (1H6.2) antibodies were used for JNK-RAPTOR PLA. Rabbit anti-mTOR (7C10) and mouse anti-RAPTOR (1H6.2) antibodies were used for mTOR-RAPTOR PLA. Mouse anti-HSF1 (E-4) and rabbit anti-AO (A11) antibodies were used for HSF1-AO PLA. Rabbit anti-AO (A11) and mouse anti-HSP60 (LK1) antibodies were used for AO-HSP60 PLA.

### Measurement of global protein translation

Live cells were incubated with 100 nM 6-FAM-dC-puromycin in complete culture medium for 30 min and analyzed by a FACSCalibur flow cytometer (BD Biosciences).

### Immunoblotting and Immunoprecipitation

Whole-cell protein extracts were prepared in cold cell-lysis buffer (100 mM NaCl, 30 mM Tris-HCl pH 7.6, 1% Triton X-100, 20 mM sodium fluoride, 1mM EDTA, 1mM sodium orthovanadate, and 1x Halt™ protease inhibitor cocktail). Proteins were transferred to nitrocellulose membranes. Following incubation with the blocking buffer (5% non-fat milk in 1x TBS-T) for 1 hour at RT, membranes were incubated with primary antibodies (1:1,000 dilution in the blocking buffer) overnight at 4 °C. After washing with 1xTBS-T for 3 times, membranes were incubated with peroxidase-conjugated secondary antibodies (1: 3,000 diluted in the blocking buffer) at RT for 1 hr. Signals were detected using SuperSignal™ West chemiluminescent substrates. For normal Co-IP, proteins were extracted using a QSonica Q125 sonicator (total process time: 15S, pulse-on time: 5S, pulse-off time: 10S, output intensity: 30%) in 1x sonication buffer (20 mM Tris, 20 mM NaCl, 1 mM EDTA, 20 mM β-glycerol-phosphate, 20 mM sodium fluoride, 4 mM sodium orthovanadate, and 1 mM DTT pH7.4, supplemented with Halt protease inhibitor cocktail). Total 1 mg whole cell lysates were incubated with A11 antibodies at 4 °C overnight. Normal rabbit IgG were used as the negative controls. The primary antibody was precipitated using Protein G magnetic beads. After washing with the lysis buffer for 3 times, beads were boiled in 1x loading buffer for 5 min before loading on SDS-PAGE. EasyBlocker (Cat#GTX425858, GeneTex) was used for blocking and anti-HSF1 (H-311) primary antibody (cat#sc-9144, Santa Cruz Biotechnology) incubation (1:1,000, O/N at 4°C). The blot was incubated with EasyBlot anti-Rabbit IgG HRP conjugated antibody (cat#GTX221666-01, GeneTex) for 1 hour. Chemiluminescent signals were captured by an iBright™ FL1000 imaging system (ThermoFisher Scientific).

### Live-Or-Dye™ Dead Cell Staining

Cells were cultured in DMEM for 48 hours before being treated with inhibitors for the required periods and collected by trypsinization. 1x10^6^ cells/ml cells were stained with Live-or-Dye™ fixable dead cell dyes (1:1,000 dilution, cat#32003, Biotium) at RT for 30 min, protected from light. Stained cells were washed once with PBS and analyzed by flow cytometry. For co-staining with cleaved caspase 3 antibody, stained cells were fixed with 4% formaldehyde (10 min, RT) and permeabilized with 0.3% Triton-X-100 in PBS (10 min, RT). The cells were stained by Cleaved Caspase 3 antibody (Asp175) (5A1E) (1:1,000, cat#9664, Cell Signaling Technology) followed by goat anti-Rabbit IgG Alexa Fluor™ 488 conjugated (1:1,000, cat#A-11034, ThermoFisher Scientific) secondary antibodies. The samples were analyzed by flow cytometry.

### Inhibitor treatments

Treatments with DTHIB, Q-VD-OPh, Liproxstatin-1, Necrostatin-1, and Wortmannin were done for 72 hours. For Congo Red and LY-2584702 treatments, cells were pretreated for 24h followed by DTHIB for an additional 72h. For treatments with NV-5138, cells were pretreated for 48 hours in a leucine-free DMEM (cat#226-024, Crystalgen) before adding DTHIB for an additional 72 hours.

### Animal studies

All animal experiments described here were approved by the institutional Animal Care and Use Committee (ACUC) of the National Cancer Institute, Frederick. Experiments on all animals were conducted following the recommendations of the Guide for the Care and Use of Laboratory Animals (National Academies Press, 2011). Animals were housed and maintained on a 12-hour light/12-hour dark cycle at temperatures between 20°C and 27°C and humidity levels of 30% to 70%, with food and water *available ad libitum*.

To establish the MPNST xenograft model, 9x10^6^ S462 MPNST cells (female) suspended with 30 % Matrigel were subcutaneously implanted the right flanks of 8-week-old female NIH-III nude mice (Charles River Laboratories). After 7 days, xenografted mice were assigned to 4 groups, eight each group, by stratified randomization based on tumor sizes (between 72 and 199 mm³). Mice were treated with KRIBB11 (65mg/kg, i.p. injection daily, cat#HY-100872, Lot#317606, Medchemexpress LLC) with and without 2% L-leucine dissolved in drinking water stored in amber-colored bottles. KRIBB11 was freshly dissolved daily in the vehicle (final concentration:10% dimethylacetamide, 50% PEG300, 20% Captisol, and 20% sterile saline).

To establish the syngeneic melanoma model, 2 × 10^6^ B16-F10-luc2 cells (male) mixed with Matrigel (1:1 ratio) in a 200μL volume were subcutaneously implanted into the right flanks of 8-week-old C57BL/6J male mice (The Jackson Laboratory). After 5 days, xenografted mice were assigned to 4 groups, eight each group, by stratified randomization based on tumor sizes (between 75 and 284 mm³). Mice were treated with KRIBB11 (65 mg/kg, i.p. injection daily, cat#T3652, Lot#132988, TargetMol Chemicals Inc.) with and without 2% L-leucine dissolved in drinking water stored in amber-colored bottles. KRIBB11 was freshly dissolved daily in the vehicle (final concentration:10% dimethylacetamide, 50% PEG300, 16% Captisol, and 24% sterile saline).

To formulate the KRIBB11 solution, KRIBB11 was first dissolved in dimethylacetamide, followed by addition of PEG300. Lastly, 40 or 50% Capitol dissolved in sterile saline was added. Tumor sizes were measured using a caliper by an animal technician blind to the study. To assess the compound’s toxicity, the body weight of tumor-bearing animals was also recorded. Tumor volumes were estimated using the formula [length (mm) × width^²^ (mm^²^)]/2.

### Quantification And Statistical Analysis

Statistical analyses were performed using Prism GraphPad 10.0 (GraphPad Software). The detailed statistical methods and sample sizes are provided in the figure legends. All results are expressed as mean±SD, mean±SEM, or median and IQR. The statistical significance is defined as: *p<0.05, **p<0.01; ***p<0.001; ****p<0.0001; n.s.: not significant. For *in vitro* experiments, sample size required was not determined a priori. The experiments were not randomized. For *in vivo* experiments, sample size required was determined by pilot experiments.

