## Supplemental Figures for "Driving proteomic imbalance in malignancy provokes proteomic catastrophe and confers tumor suppression"

**LIST OF SUPPLEMENTARY MATERIALS**

The supplementary materials include 7 supplementary figures and their legends.

**Figure S1**: **HSF1 is required to suppress proteomic instability and amyloidogenesis in MPNST cells, related to Figure 1.**

**Figure S2: HSF1 counters the tumor-suppressive effects of amyloids in MPNST cells, related to Figure 2.**

**Figure S3: JNK senses *HSF1* deficiency to repress mTORC1 and protein translation in MPNST cells, related to Figure 3.**

**Figure S4: Protein translation is causally related to amyloidogenesis in MPNST cells, related to Figure 4.**

**Figure S5: mTORC1 stimulation aggravates the amyloidogenesis and cytotoxicity elicited by HSF1 inhibition, related to Figure 5.**

**Figure S6: Concurrent HSF1 inhibition and mTORC1 stimulation instigates widespread non-apoptotic cell death, related to Figure 6.**

**Figure S7: Concurrent HSF1 inhibition and mTORC1 stimulation suppress *in vivo* tumor growth, related to Figure 7.**

### Figure S1

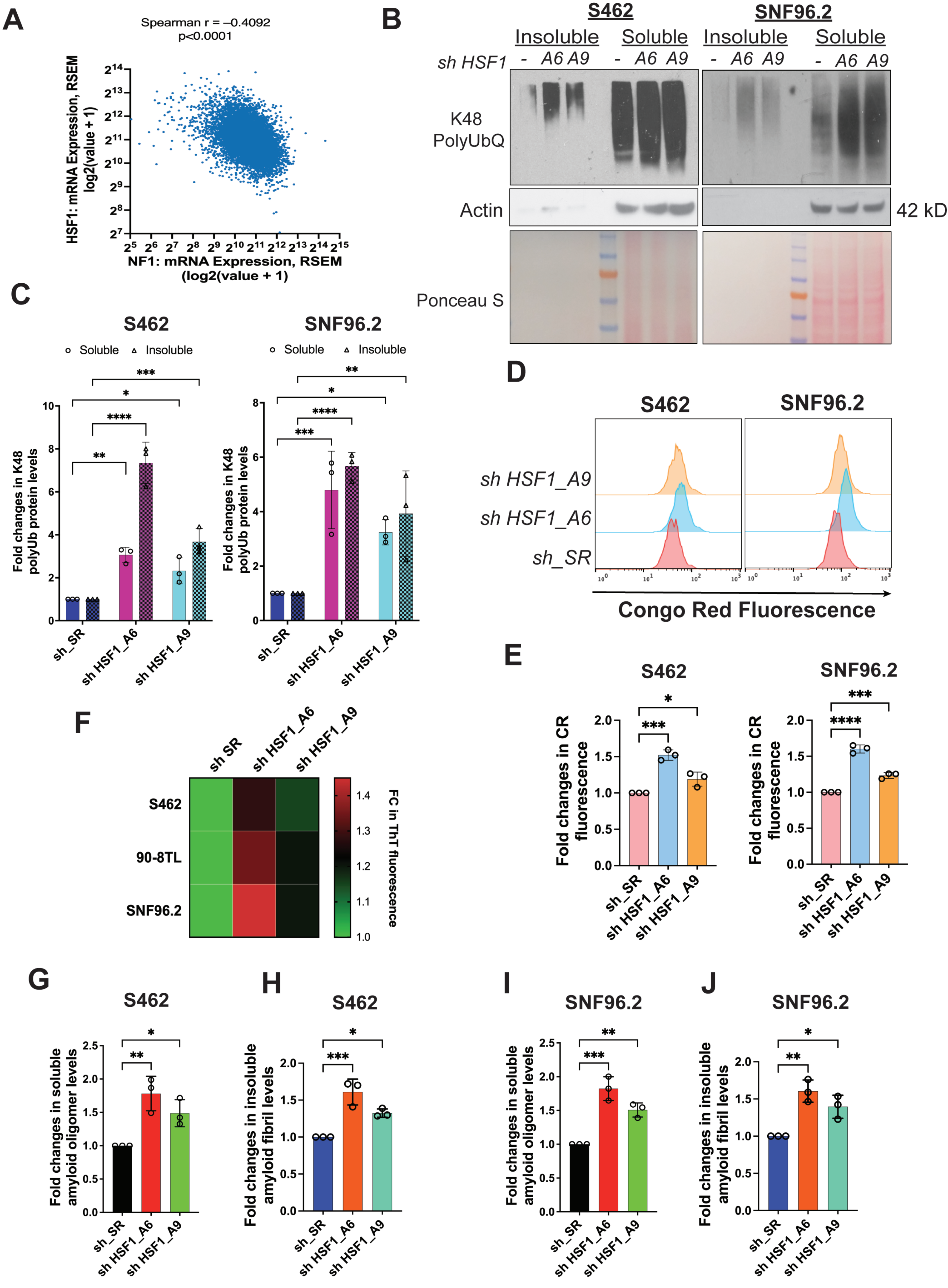
A
Spearman r = –0.4092
p<0.0001
HSF1: mRNA Expression, RSEM log2(value + 1)
NF1: mRNA Expression, RSEM (log2(value + 1)
B
S462
SNF96.2
Insoluble
Soluble
sh HSF1
A6
A9
K48 PolyUbQ
Actin
42 kD
Ponceau S
C
S462
SNF96.2
Soluble
Insoluble
Fold changes in K48 polyUb protein levels
sh_SR
sh HSF1_A6
sh HSF1_A9
D
sh HSF1_A9
sh HSF1_A6
sh_SR
Congo Red Fluorescence
E
Fold changes in CR fluorescence
F
sh SR
sh HSF1_A6
sh HSF1_A9
S462
90-8TL
SNF96.2
FC in ThT fluorescence
G
Fold changes in soluble amyloid oligomer levels
H
Fold changes in insoluble amyloid fibril levels
I
J

#### SUPPLEMENTARY FIGURE LEGENDS

**Figure S1: HSF1 is required to suppress proteomic instability and amyloidogenesis in MPNST cells.**

(A) Reverse correlation between *NF1* and *HSF1* mRNA levels in human cancer tissues (n=10,071, Spearman's correlation). Data are generated by the TGCA Research Network (https://www.cancer.gov/tcga). (B) The impacts of *HSF1* depletion on protein polyubiquitination in S462 cells and SNF96.2 cells. Cells were transduced with lentiviral shRNAs for 4 days and global protein Lys48-specific ubiquitination was detected by immunoblotting in both detergent-soluble and -insoluble fractions of whole cell lysates. (C) Protein ubiquitination was quantitated using Fiji imaging software (B) (mean ± SD, n=3 independent experiments, Two-way ANOVA). (D) Congo Red (CR) staining of S462 and SNF96.2 cells with and without *HSF1* depletion, analyzed by flow cytometry. (E) Quantitation of CR staining in (D) using the median fluorescence intensity (mean ± SD, n=3 independent experiments, One-way ANOVA). (F) Thioflavin T (ThT) staining of S462, 90-8TL, and SNF96.2 cells with and without *HSF1* depletion, analyzed by flow cytometry. The median fluorescence intensity was presented as a heatmap. (G)-(J) Quantitation of soluble amyloid oligomers and insoluble amyloid fibrils in S462 and SNF96.2 cells by ELISA using A11 and OC antibodies, respectively (mean ± SD, n=3 independent experiments, One-way ANOVA).

### Figure S2

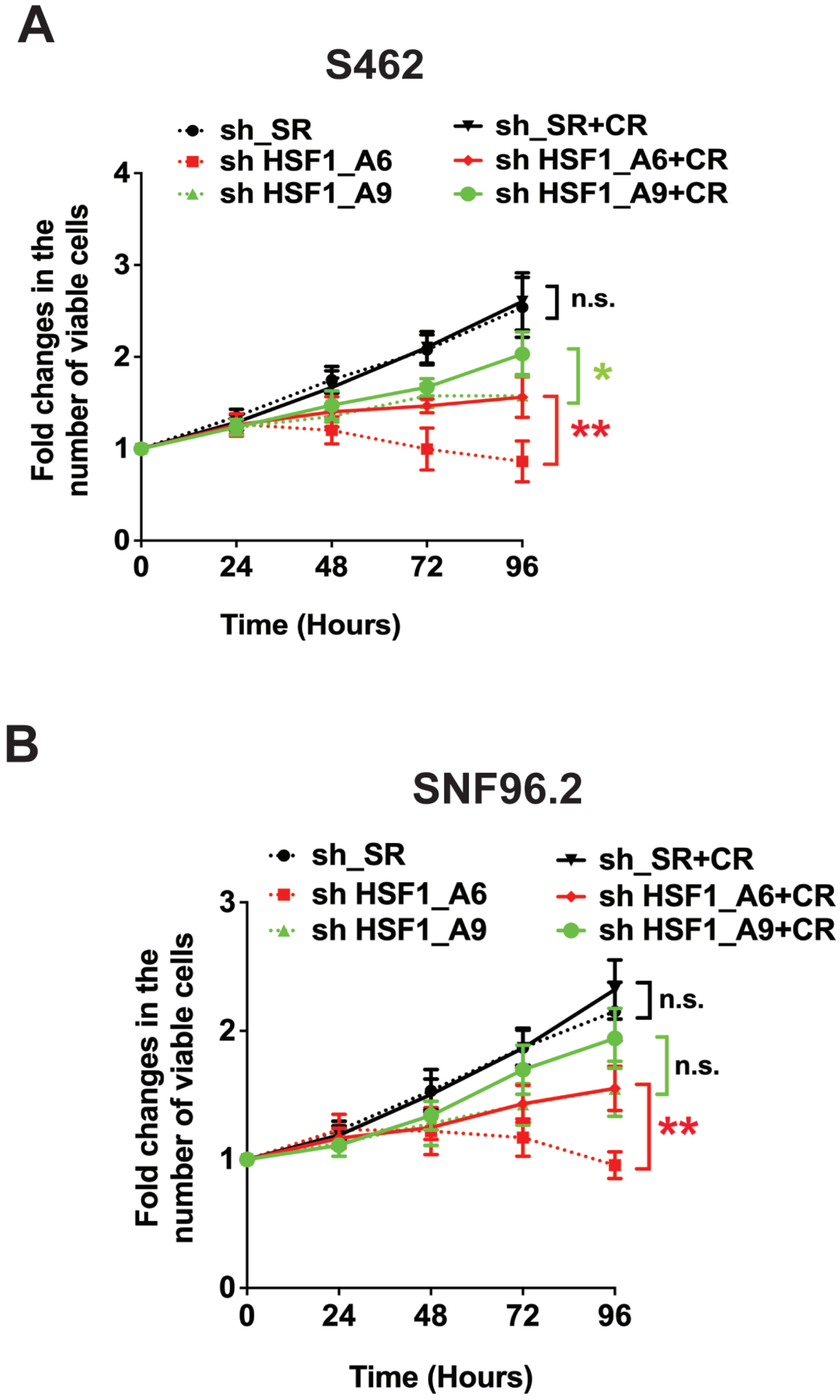
A
S462
sh_SR
sh HSF1_A6
sh HSF1_A9
sh_SR+CR
sh HSF1_A6+CR
sh HSF1_A9+CR
Fold changes in the number of viable cells
0
1
2
3
4
0
24
48
72
96
Time (Hours)
n.s.
*
**
B
SNF96.2
sh_SR
sh HSF1_A6
sh HSF1_A9
sh_SR+CR
sh HSF1_A6+CR
sh HSF1_A9+CR
Fold changes in the number of viable cells
0
1
2
3
0
24
48
72
96
Time (Hours)
n.s.
n.s.
**

**Figure S2: HSF1 counters the tumor-suppressive effects of amyloids in MPNST cells.**

(A) The growth curves of S462 cells with and without *HSF1* depletion and CR treatment (mean ± SD, n=3 independent experiments, Two-way ANOVA). (B) The growth curves of SNF96.2 cells with and without *HSF1* depletion and CR treatment (mean ± SD, n=3 independent experiments, Two-way ANOVA).

### Figure S3

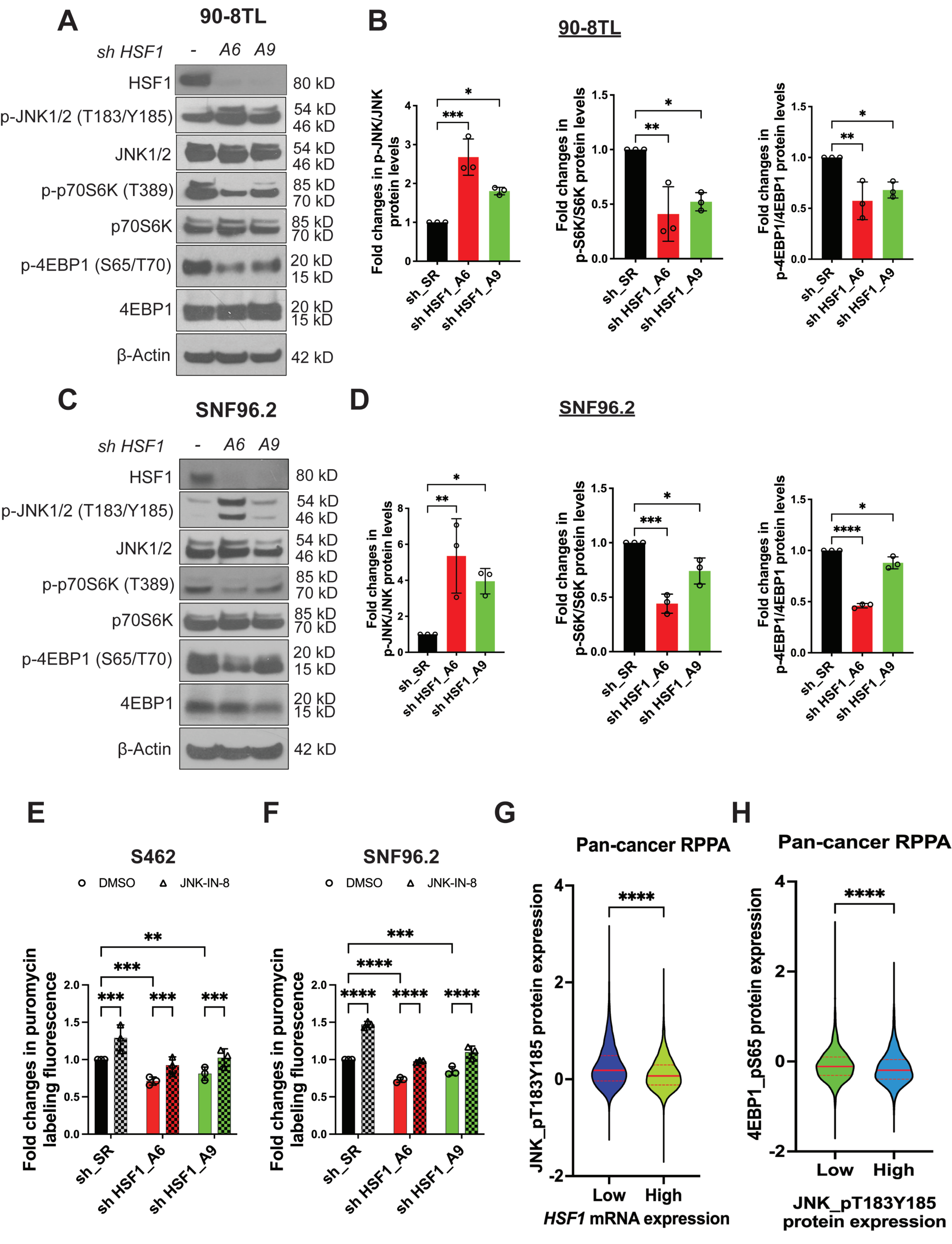
A
90-8TL
sh HSF1
-
A6
A9
HSF1
80 kD
p-JNK1/2 (T183/Y185)
54 kD
46 kD
JNK1/2
54 kD
46 kD
p-p70S6K (T389)
85 kD
70 kD
p70S6K
85 kD
70 kD
p-4EBP1 (S65/T70)
20 kD
15 kD
4EBP1
20 kD
15 kD
β-Actin
42 kD
B
90-8TL
Fold changes in p-JNK/JNK protein levels
Fold changes in p-S6K/S6K protein levels
Fold changes in p-4EBP1/4EBP1 protein levels
sh_SR
sh HSF1_A6
sh HSF1_A9
C
SNF96.2
sh HSF1
-
A6
A9
HSF1
80 kD
p-JNK1/2 (T183/Y185)
54 kD
46 kD
JNK1/2
54 kD
46 kD
p-p70S6K (T389)
85 kD
70 kD
p70S6K
85 kD
70 kD
p-4EBP1 (S65/T70)
20 kD
15 kD
4EBP1
20 kD
15 kD
β-Actin
42 kD
D
SNF96.2
Fold changes in p-JNK/JNK protein levels
Fold changes in p-S6K/S6K protein levels
Fold changes in p-4EBP1/4EBP1 protein levels
sh_SR
sh HSF1_A6
sh HSF1_A9
E
S462
DMSO
JNK-IN-8
Fold changes in puromycin labeling fluorescence
sh_SR
sh HSF1_A6
sh HSF1_A9
F
SNF96.2
DMSO
JNK-IN-8
Fold changes in puromycin labeling fluorescence
sh_SR
sh HSF1_A6
sh HSF1_A9
G
Pan-cancer RPPA
JNK_pT183Y185 protein expression
Low
High
HSF1 mRNA expression
H
Pan-cancer RPPA
4EBP1_pS65 protein expression
Low
High
JNK_pT183Y185 protein expression

**Figure S3: JNK senses *HSF1* deficiency to repress mTORC1 and protein translation in MPNST cells.**

(A) Detection of JNK activation and mTORC1 inhibition following *HSF1* depletion in 90-8TL cells by immunoblotting. (B) Quantitation of (A) using Fiji imaging software (mean ± SD, n=3 independent experiments, One-way ANOVA). (C) Detection of JNK activation and mTORC1 inhibition following *HSF1* depletion in SNF96.2 cells by immunoblotting. (D) Quantitation of (C) using Fiji imaging software (mean ± SD, n=3 independent experiments, One-way ANOVA). (E) and (F) Measurement of protein translation rates by puromycin labeling in S462 and SNF96.2 cells with and without *HSF1* depletion and JNK inhibition (mean ± SD, n=3 independent experiments, Two-way ANOVA). (G) Violin plots of JNK T183/Y185 phosphorylation in human cancer tissues with low and high *HSF1* expression (median and IQR, n=3,526 or 3,527, Mann-Whitney U test). Data are generated by the TGCA Research Network (https://www.cancer.gov/tcga) and the Cancer Proteome Atlas (https://tcpaportal.org/tcpa/). (H) Violin plots of 4EBP1 S65 phosphorylation in human cancer tissues with low and high JNK T183/Y185 phosphorylation (median and IQR, n=3,526 or 3,527, Mann-Whitney U test). Data are generated by the Cancer Proteome Atlas (https://tcpaportal.org/tcpa/).

### Figure S4

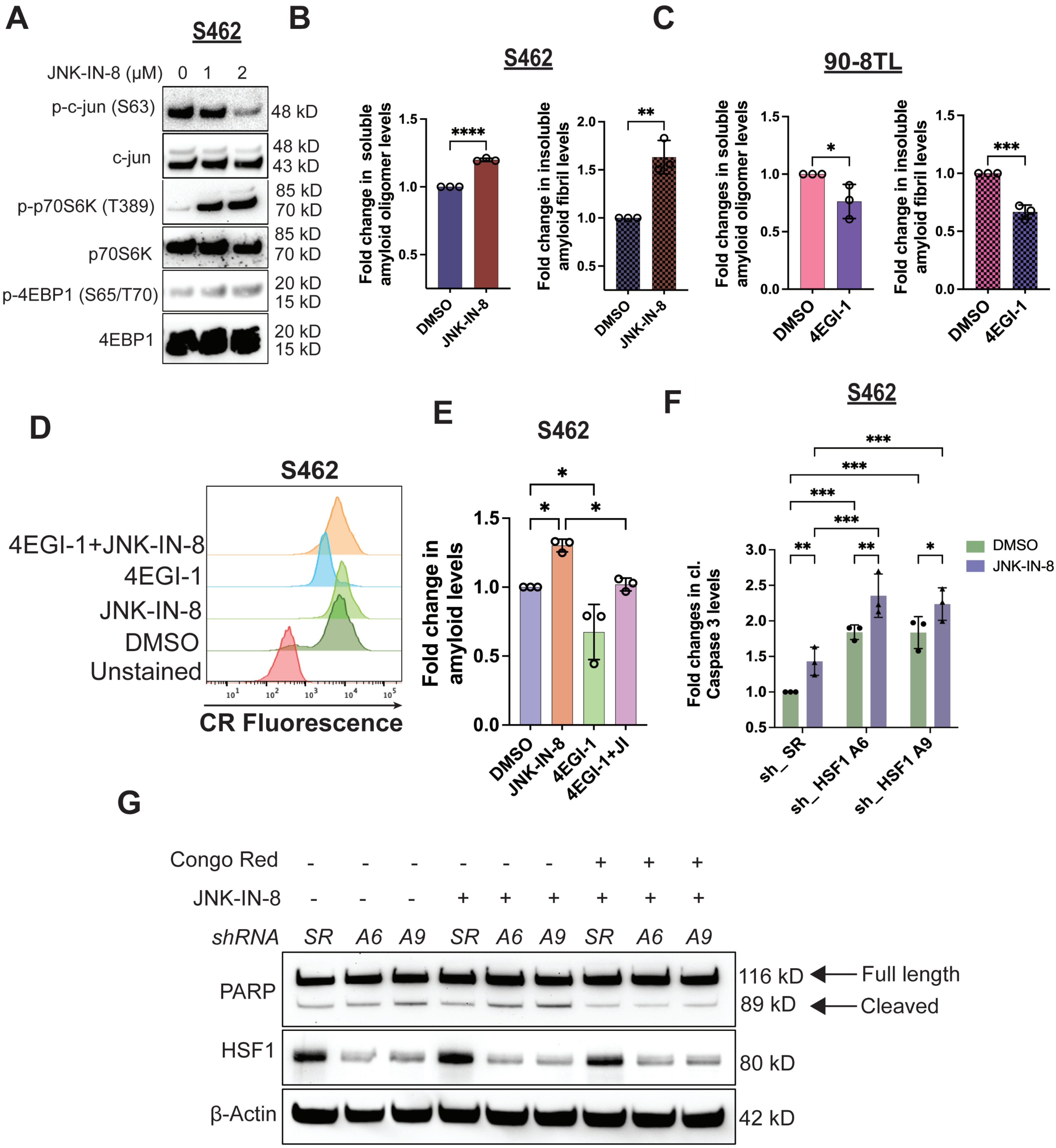
A
S462
JNK-IN-8 (µM) 0 1 2
p-c-jun (S63)
48 kD
c-jun
48 kD
43 kD
p-p70S6K (T389)
85 kD
70 kD
p70S6K
85 kD
70 kD
p-4EBP1 (S65/T70)
20 kD
15 kD
4EBP1
20 kD
15 kD
B
S462
Fold change in soluble amyloid oligomer levels
Fold change in insoluble amyloid fibril levels
DMSO
JNK-IN-8
C
90-8TL
Fold changes in soluble amyloid oligomer levels
Fold change in insoluble amyloid fibril levels
DMSO
4EGI-1
D
S462
4EGI-1+JNK-IN-8
4EGI-1
JNK-IN-8
DMSO
Unstained
CR Fluorescence
E
S462
Fold change in amyloid levels
DMSO
JNK-IN-8
4EGI-1
4EGI-1+JI
F
S462
Fold changes in cl. Caspase 3 levels
DMSO
JNK-IN-8
sh_ SR
sh_ HSF1 A6
sh_ HSF1 A9
G
Congo Red
JNK-IN-8
shRNA
SR A6 A9 SR A6 A9 SR A6 A9
PARP
116 kD
Full length
89 kD
Cleaved
HSF1
80 kD
β-Actin
42 kD

**Figure S4: Protein translation is causally related to amyloidogenesis in MPNST cells.**

(A) Detection of mTORC1 stimulation following JNK inhibition in S462 cells by immunoblotting. Cells were treated with JNK-IN-8 for 2 days. (B) Quantitation of soluble AOs and insoluble AFs by ELISA in S462 cells with and without 3 μM JNK-IN-8 treatment for 3 days (mean ± SD, n=3 independent experiments, Student's t test). (C) Quantitation of soluble AOs and insoluble AFs by ELISA in 90-8TL cells with and without 50 μM 4EGI-1 treatment for 3 days (mean ± SD, n=3 independent experiments, Student's t test). (D) CR staining of S462 cells with and without JNK-IN-8 and 4EGI-1 treatment, detected by flow cytometry. (E) Quantitation of (D) using the median fluorescence intensity (mean ± SD, n=3 independent experiments, One-way ANOVA). (F) Quantitation of caspase 3 cleavage by ELISA in S462 cells with and without *HSF1* depletion and 3 μM JNK-IN-8 treatment (mean ± SD, n=3 independent experiments, Two-way ANOVA). (G) Immunoblotting detection of PARP cleavage in 90-8TL cells treated with and without 3μM JNK-IN-8 alone or combined JNK-IN-8 and 20 μM CR following *HSF1* depletion.

### Figure S5

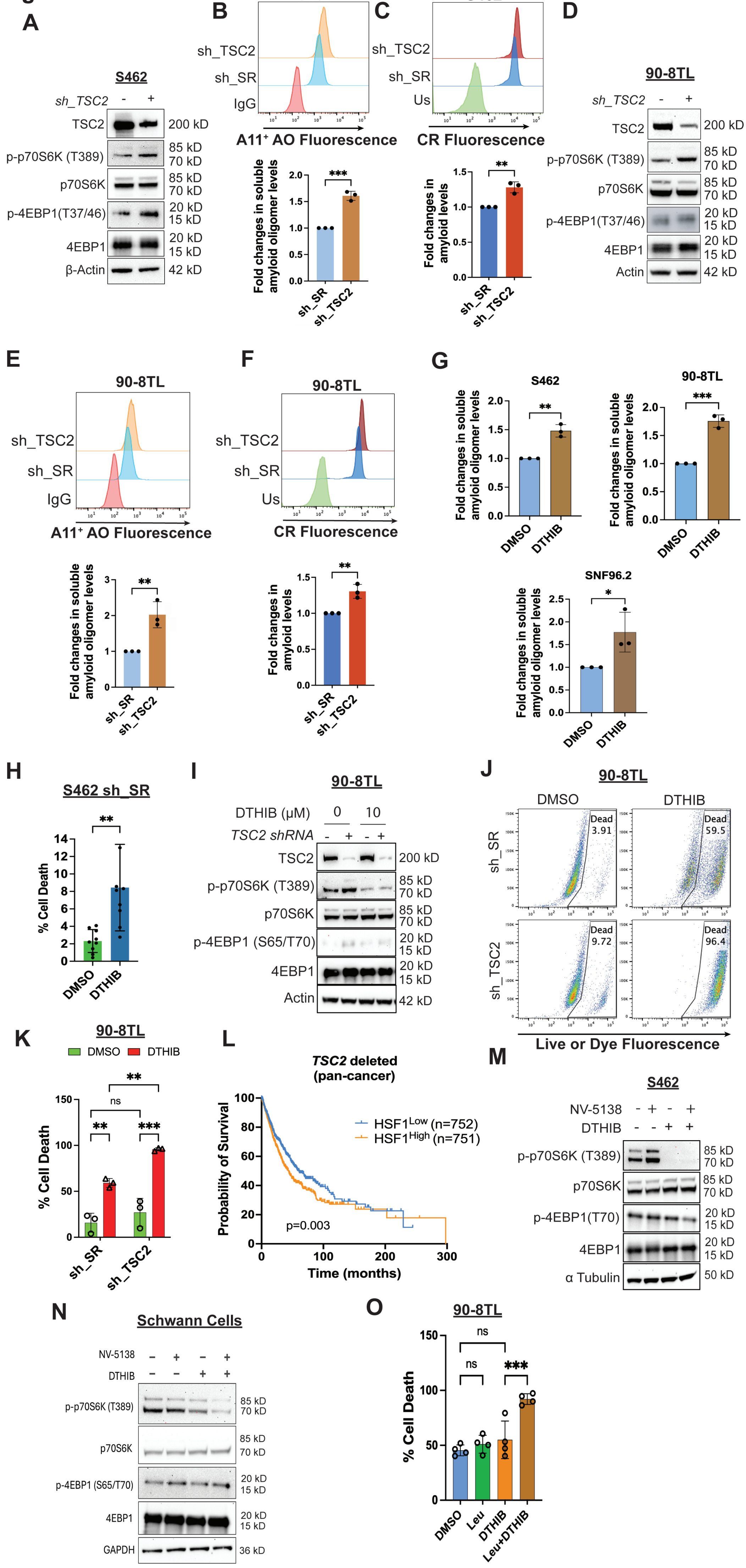

A
S462
sh_TSC2 - +
TSC2 200 kD
p-p70S6K (T389) 85 kD 70 kD
p70S6K 85 kD 70 kD
p-4EBP1(T37/46) 20 kD 15 kD
4EBP1 20 kD 15 kD
β-Actin 42 kD
B
S462
sh_TSC2
sh_SR
IgG
A11+ AO Fluorescence
Fold changes in soluble amyloid oligomer levels
***
sh_SR
sh_TSC2
C
S462
sh_TSC2
sh_SR
Us
CR Fluorescence
Fold changes in amyloid levels
**
D
90-8TL
sh_TSC2 - +
TSC2 200 kD
p-p70S6K (T389) 85 kD 70 kD
p70S6K 85 kD 70 kD
p-4EBP1(T37/46) 20 kD 15 kD
4EBP1 20 kD 15 kD
Actin 42 kD
E
90-8TL
A11+ AO Fluorescence
F
90-8TL
CR Fluorescence
G
S462
90-8TL
SNF96.2
DMSO
DTHIB
H
S462 sh_SR
% Cell Death
I
90-8TL
DTHIB (μM) 0 10
TSC2 shRNA - + - +
TSC2 200 kD
p-p70S6K (T389) 85 kD 70 kD
p70S6K 85 kD 70 kD
p-4EBP1 (S65/T70) 20 kD 15 kD
4EBP1 20 kD 15 kD
Actin 42 kD
J
90-8TL
DMSO
DTHIB
Dead 3.91
Dead 59.5
Dead 9.72
Dead 96.4
Live or Dye Fluorescence
K
90-8TL
DMSO DTHIB
ns
L
TSC2 deleted (pan-cancer)
HSF1Low (n=752)
HSF1High (n=751)
Probability of Survival
Time (months)
p=0.003
M
S462
NV-5138 - + - +
DTHIB - - + +
p-p70S6K (T389) 85 kD 70 kD
p70S6K 85 kD 70 kD
p-4EBP1(T70) 20 kD 15 kD
4EBP1 20 kD 15 kD
α Tubulin 50 kD
N
Schwann Cells
NV-5138 - + - +
DTHIB - - + +
p-p70S6K (T389) 85 kD 70 kD
p70S6K 85 kD 70 kD
p-4EBP1 (S65/T70) 20 kD 15 kD
4EBP1 20 kD 15 kD
GAPDH 36 kD
O
90-8TL
DMSO
Leu
DTHIB
Leu+DTHIB

**Figure S5: mTORC1 stimulation aggravates the amyloidogenesis and cytotoxicity elicited by HSF1 inhibition.**

(A) Immunoblotting detection of TSC2 expression and mTORC1 activity in S462 cells following stable *TSC2* knockdown. (B) and (C) Quantitation of amyloid levels in S462 cells following stable *TSC2* knockdown by either A11 antibody or CR staining, analyzed by flow cytometry (mean ± SD, n=3 independent experiments, Student's t test). (D) Immunoblotting detection of TSC2 expression and mTORC1 activity in 90-8TL cells following stable *TSC2* knockdown. (E) and (F) Quantitation of amyloid levels in 90-8TL cells following stable *TSC2* knockdown by either A11 antibody or CR staining, analyzed by flow cytometry (mean ± SD, n=3 independent experiments, Student's t test). (G) Quantitation of soluble amyloid oligomers by ELISA in S462, 90-8TL, and SNF96.2 cells following 10 µM DTHIB treatment for 3 days (mean ± SD, n=3 independent experiments, Student's t test). (H) Quantitation of cytotoxicity using Live-or-Dye staining in SR control S462 cells with and without 10 µM DTHIB treatment for 3 days (mean ± SD, n=9 independent experiments, Student's t test). (I) Immunoblotting detection of TSC2 expression and mTORC1 activity in 90-8TL cells following stable *TSC2* knockdown. (J) and (K) Quantitation of cytotoxicity using Live-or-Dye staining in 90-8TL cells with and without stable *TSC2* knockdown and 10 µM DTHIB treatment for 3 days (mean ± SD, n=3 independent experiments, Two-way ANOVA). (L) Kaplan-Meier survival curves of patients whose tumors harbor *TSC2* deletion, stratified by *HSF1* mRNA expression (n=752 or 751, Log-rank test). Data are generated by the TGCA Research Network (https://www.cancer.gov/tcga). (M) Immunoblotting detection of mTORC1 stimulation by 500 µM NV-5138 in S462 cells with and without 10 µM DTHIB treatment. (N) Immunoblotting detection of mTORC1 stimulation by 500 µM NV-5138 in immortalized human Schwann cells with and without 10 µM DTHIB treatment.

(O) Quantitation of cytotoxicity by flow cytometry using Live-or-Dye stains 90-8TL cells with and without 500 μM L-leucine stimulation and 10 μM DTHIB treatment (mean ± SD, n=4 independent experiments, One-way ANOVA).

### Figure S6

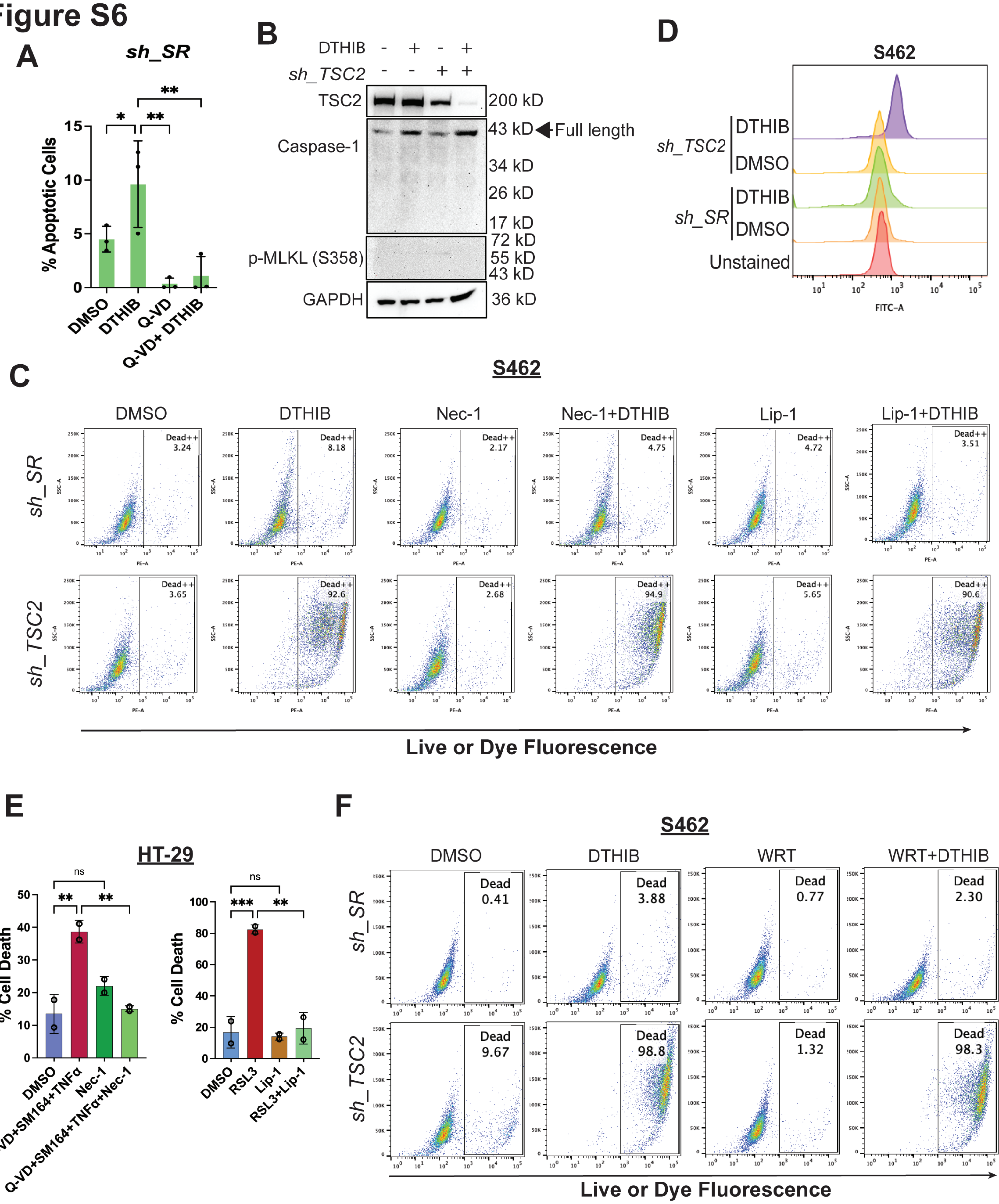
A
sh_SR
% Apoptotic Cells
DMSO
DTHIB
Q-VD
Q-VD+ DTHIB
B
DTHIB
sh_TSC2
TSC2
200 kD
Caspase-1
43 kD
Full length
34 kD
26 kD
17 kD
p-MLKL (S358)
72 kD
55 kD
43 kD
GAPDH
36 kD
D
S462
sh_TSC2
DTHIB
DMSO
sh_SR
DTHIB
DMSO
Unstained
FITC-A
C
S462
DMSO
DTHIB
Nec-1
Nec-1+DTHIB
Lip-1
Lip-1+DTHIB
sh_SR
sh_TSC2
Dead++
3.24
8.18
2.17
4.75
4.72
3.51
3.65
92.6
2.68
94.9
5.65
90.6
SSC-A
PE-A
Live or Dye Fluorescence
E
HT-29
% Cell Death
DMSO
Q-VD+SM164+TNFα
Nec-1
Q-VD+SM164+TNFα+Nec-1
RSL3
Lip-1
RSL3+Lip-1
ns
F
S462
DMSO
DTHIB
WRT
WRT+DTHIB
sh_SR
sh_TSC2
Dead
0.41
3.88
0.77
2.30
9.67
98.8
1.32
98.3
Live or Dye Fluorescence

**Figure S6: Concurrent HSF1 inhibition and mTORC1 stimulation instigates widespread necrotic cell death.**

(A) Quantitation of apoptosis, defined by caspase 3 cleavage, in SR control S462 cells treated with and without 10 μM DTHIB alone or combined DTHIB and 30 μM Q-VD-OPh for 3 days (mean ± SD, n=3 independent experiments, One-way ANOVA). (B) Immunoblotting detection of caspase 1 cleavage and MLKL Ser358 phosphorylation in *TSC2*-deficient S462 cells treated with and without 10 μM DTHIB. (C) Representative scatter plots of Live-or-Dye staining of S462 cells treated with DTHIB alone or combined DTHIB and either Necrostatin-1 (Nec-1) or liproxstatin-1 (Lip-1). Quantitation is presented in Figure 6C. (D) Measurement of lipid peroxides using Liperfluo staining in *TSC2*-deficient S462 cells treated with and without 10 μM DTHIB, analyzed by flow cytometry. (E) 30 μM Necrostatin-1 and 20 μM Liproxstatin-1 effectively blocked necroptosis and ferroptosis induced in HT-29 cells, respectively. Cell death was measured by Live-or-Dye staining (mean ± SD, n=2 independent experiments, One-way ANOVA). (F) Representative scatter plots of Live-or-Dye staining of S462 cells treated with DTHIB alone or combined DTHIB and wortmannin (WRT). Quantitation is presented in Figure 6F.

Figure S7

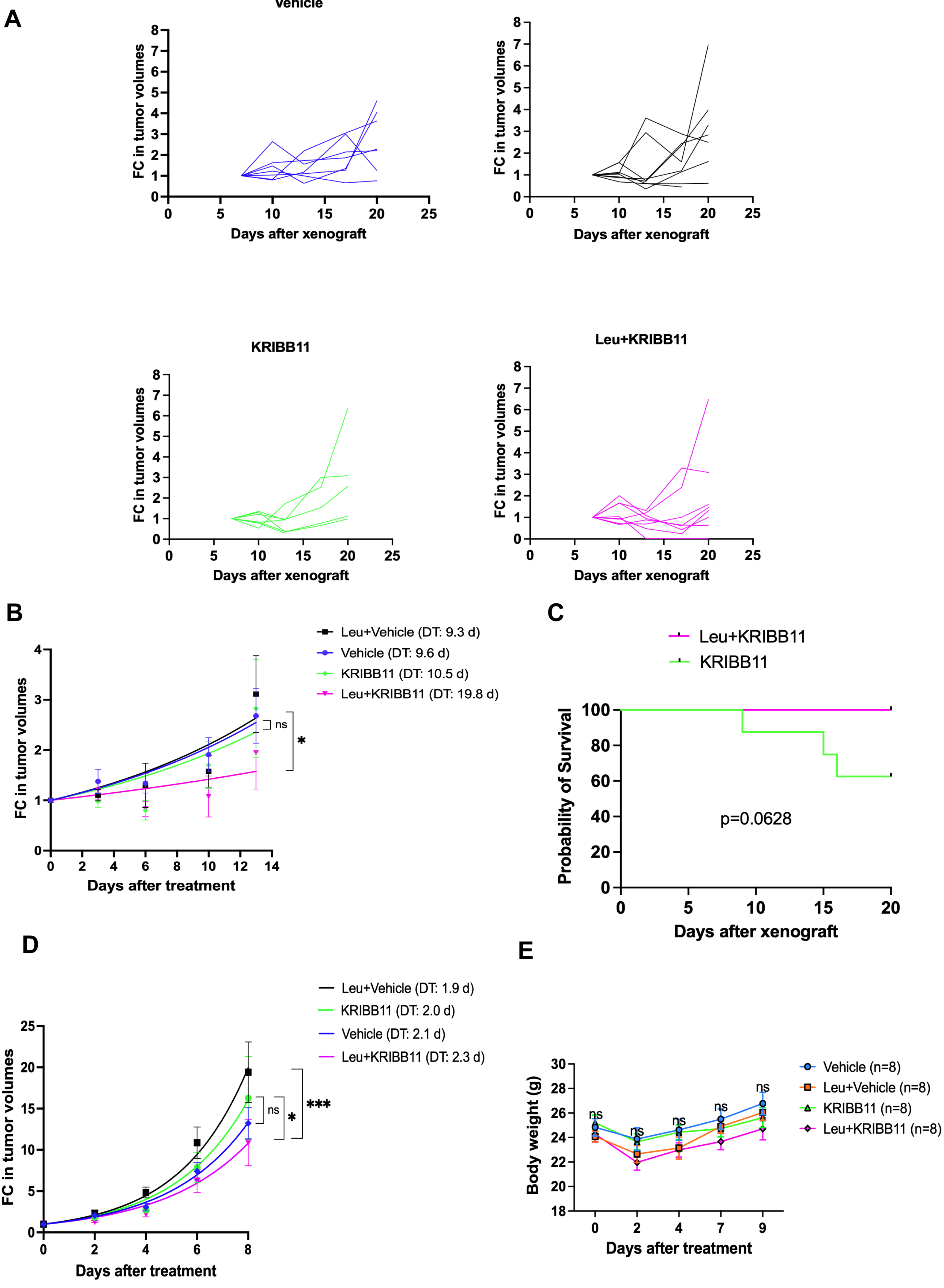

A
Vehicle
Leu
KRIBB11
Leu+KRIBB11
FC in tumor volumes
Days after xenograft
B
Leu+Vehicle (DT: 9.3 d)
Vehicle (DT: 9.6 d)
KRIBB11 (DT: 10.5 d)
Leu+KRIBB11 (DT: 19.8 d)
ns
*
FC in tumor volumes
Days after treatment
C
Leu+KRIBB11
KRIBB11
Probability of Survival
p=0.0628
Days after xenograft
D
Leu+Vehicle (DT: 1.9 d)
KRIBB11 (DT: 2.0 d)
Vehicle (DT: 2.1 d)
Leu+KRIBB11 (DT: 2.3 d)
ns
*
***
FC in tumor volumes
Days after treatment
E
Vehicle (n=8)
Leu+Vehicle (n=8)
KRIBB11 (n=8)
Leu+KRIBB11 (n=8)
ns
Body weight (g)
Days after treatment

**Figure S7: Concurrent HSF1 inhibition and mTORC1 stimulation suppresses in vivo tumor growth.**

(A) Individual growth curves of xenografted human S462 MPNSTs in NIH-III nude mice. (B) FC in tumor volumes of xenografted human S462 MPNSTs are fitted to the exponential growth equation to derive the tumor volume doubling time (DT) and exponential growth constants. Growth constants are compared among curves (mean ± SEM, n=8 mice per group, One-way ANOVA). (C) Kaplan-Meier survival curves of NIH-III nude mice treated with either KRIBB11 alone or combined KRIBB11 with L-leucine (logrank test). (D) FC in tumor volumes of xenografted murine B16F10 melanomas are fitted to the exponential growth equation to derive the tumor volume doubling time (DT) and exponential growth constants. Growth constants are compared among curves (mean ± SEM, n=8 mice per group, One-way ANOVA). (E) Changes in body weights of C57BL/6J mice during treatments (mean ± SD, n=8 mice per group, Two-way ANOVA).
